# Myt Transcription Factors prevent stress-response gene over-activation to enable postnatal pancreatic β cell proliferation and function

**DOI:** 10.1101/773846

**Authors:** Ruiying Hu, Emily Walker, Yanwen Xu, Chen Huang, Chen Weng, Gillian E. Erickson, Anastasia Golovin, Xiaodun Yang, Marcella Brissova, Appakalai N. Balamurugan, Christopher V. E. Wright, Yan Li, Roland Stein, Guoqiang Gu

**Author notes:** These authors contributed equally to the work (Ruiying Hu, Emily Walker, Yanwen Xu, Chen Huang). Corresponding authors and lead contact: Guoqiang Gu.

## Abstract

Although stress response maintains cell function and survival under adverse conditions, over-activation of late-stage stress-gene effectors causes dysfunction and death. Here we show that the Myelin Transcription Factors (Myt 1, 2, and 3 TFs) prevent this over-activation. Co-inactivating *Myt TFs* in mouse pancreatic progenitors compromised postnatal β-cell function, proliferation, and survival, preceded by upregulation of late-stage stress-response genes *Activating Transcription Factors* (e.g., *Atf4*) and *Heat Shock Proteins* (*Hsps*). Myt1 binds the putative enhancers of *Atf4* and *Hsp*s, whose over-expression in mouse β cells largely recapitulated the *Myt* mutant phenotypes. Moreover, Myt(MYT)-TF levels were upregulated in functional mouse and human β cells by metabolic stress but downregulated in those of type 2 diabetic islets that display *ATF4* and *HSP* over-activation. Lastly, human *MYT* knockdown caused stress-gene over-activation and death in Endo-βH1 cells. These findings suggest that the Myt TFs restrict stress-response to physiologically tolerable levels in mice and human.

## Introduction

The unfolded-protein response (UPR) and oxidative stress response (OSR) are central protectors of cell function and survival (Juliana et al., 2017; Sies et al., 2017). Unfolded proteins in the endoplasmic reticulum (ER) sequester resident chaperones such as GRP78/Bip and stimulate early sensor-effectors of the stress response, including Inositol Requiring 1 (IRE1), Protein Kinase RNA-like Endoplasmic Reticulum Kinase (PERK), and Activating Transcription Factor 6 (ATF6) (Cao and Kaufman, 2014). Their activation downregulates the overall protein translation but preferentially induces the production of several later-acting stress-pathway effectors such as Atf4, heat-shock proteins (Hsps), proteins required for endoplasmic-reticulum-associated protein degradation (ERAD) via splicing, transcription, translation, and/or post-translational mechanisms (Papa, 2012; Shen et al., 2002). These responses serve to facilitate nascent protein folding and to reduce the proportion of mis-folded proteins. Moreover, conditions causing cytoplasmic protein denaturation, such as heat shock and reactive-oxygen species (ROS), activate heat-shock factors (HSFs) that induce transcription of *Hsp* and chaperonin (*Cct*) genes (Fulda et al., 2010; Ostling et al., 2007). ROS also activates the Nrf2 transcription factor (TF) to induce the production of enzymes that remove toxic reactive-oxygen species (Sies et al., 2017). Collectively, these responses allow cells to survive in stressful environments. However, chronic and/or over-activation of the stress response causes cell dysfunction and death (Hotamisligil and Davis, 2016). For cells such as neuronal and pancreatic islet β cells that cannot be readily regenerated, a substantial loss of cell numbers in this manner could permanently compromise organ function and cause neuronal diseases and type 2 diabetes (T2D) (Cao and Kaufman, 2014; Chaari, 2019).

Over a long lifespan, pancreatic β cells make exceedingly large amounts of insulin, representing roughly 5–10% of the total protein pool, to sustain the ability for cyclical insulin secretion as the principal factor that maintains glucose homeostasis. This production level inevitably produces some mis-folded proinsulin, sometimes up to 20% of nascent proinsulin, in the ER – sufficiently high to induce UPR (Cao and Kaufman, 2014; Sun et al., 2015; Szabat et al., 2016). Increased mitochondrial glucose metabolism in β cells is not only the main inducer for insulin secretion, but also increases ROS production. While low ROS levels promote insulin secretion, high levels activate OSR and UPR (Guo et al., 2013; Hartl et al., 2011). The UPR and OSR of active, healthy β cells facilitates protein homeostasis and promotes proper insulin secretion and β-cell proliferation (Lipson et al., 2006; Sharma et al., 2015; Szabat et al., 2016). However, sustained high levels of stress signals attenuate general protein translation and contribute to β-cell dysfunction by activating CHOP and other pro-apoptotic genes (Iurlaro and Munoz-Pinedo, 2016). Consequently, β cells are challenged to tune their stress response within a range that allows robust secretory function to be balanced with healthy cell survival (Fonseca et al., 2011; Song et al., 2008).

The *Myelin Transcription Factors* (*Myt TFs*) [i.e. *Myt1* (*Nzf2*), *Myt2* (*Myt1L* or *Nzf1*), and *Myt3* (*Nzf3* or *St18*)] are responsive to stress signals. For example, hyperglycemia and hyperlipidemia potentiate *Myt3* expression in β cells (Henry et al., 2014; Tennant et al., 2012), a response associated either with cell survival (Henry et al., 2014) or cell death (Tennant et al., 2016). This family of zinc-finger proteins is also expressed in neural and neuro-endocrine cells (Gu et al., 2004; Henry et al., 2014; Kim and Hudson, 1992; Tennant et al., 2012; Vasconcelos et al., 2016; Wang et al., 2007), wherein they regulate cell differentiation by repressing expression of non-neuronal genes (Vasconcelos et al., 2016). Precisely defining Myt1, 2 and/or 3 activities, however, has been greatly hampered by the genetic compensation between the members (Wang et al., 2007).

Myt1, 2, and 3 were recently shown to be largely dispensable for β-cell differentiation (Liu et al., 2019). Yet they were found to repress *Synaptotagmin 4* (*Syt4*) expression in neonatal islet β cells, a key regulator of vesicle–plasma membrane fusion and insulin secretion in adult-stage β cells (Huang et al., 2018). Normally, Syt4 levels increase during postnatal β-cell maturation and are required for mediating glucose-sensitive insulin release. Early embryonic removal of *Myt1*, *2*, and *3* by expression of *Pdx1*-driven Cre in *Myt1^F/F^; Myt2^F/F^; Myt3^F/F^* mice (termed *6F; Pdx1^Cre^* hereafter) resulted in defective insulin secretion in post-weaning islets due, in part, to the precocious activation of Syt4 production. Yet overexpression of *Syt4* in perinatal β cells did not result in overt diabetes, thus leading to the conclusion that it alone cannot be responsible for all the effects observed in *6F; Pdx1^Cre^* mice (Huang et al., 2018), and suggesting that other cellular processes must be dysregulated in the absence of Myt TFs. We show here their importance in controlling an appropriate level of activity of cellular stress pathways, which is very influential in maintaining normal β-cell function, proliferation, and survival.

## Results

### Inactivation of *Myt* genes compromised β-cell proliferation and survival

To gain clues regarding the processes regulated by Myt TFs in β cells, the physiological and cellular defects of *6F; Pdx1^Cre^* (i.e., *Myt1^F/F^; Myt2^F/F^; Myt3^F/F^; Pdx1^Cre^*) mice were examined in greater detail. Myt proteins were efficiently inactivated in *6F; Pdx1^Cre^* pancreatic cells (Figure S1A1-A6). There was no change in Myt levels in the hypothalamus, where some spurious *Pdx1^Cre^* activity was reported (Figure S1B0-B8) (Schwartz et al., 2010). There were also no obvious defects in the exocrine pancreas of *6F; Pdx1^Cre^* mice (Figure S1C1-C2), consistent with the endocrine-specific pattern of Myt TF production (Figure S1A1-A3).

*6F; Pdx1^Cre^* mutant mice displayed a higher fasting blood-glucose level as early as postnatal day 14 (P14) in comparison to the wild-type, *Pdx1^Cre^*, and *Myt1^F/F^; Myt2^F/F^; Myt3^F/F^* (termed *6F*) controls, and this difference became more pronounced as the mice aged (Figure 1A, Table S1). This temporal pattern correlated with reductions in both *6F; Pdx1^Cre^* body weight (Figure 1A) and glucose-induced plasma insulin level (Figure 1B). There was no change in the plasma glucagon level (Figure 1C), an α-cell–secreted hormone that induces gluconeogenesis in (for example) the liver, nor a detectably compromised insulin sensitivity (Figure 1D). Ruling out whole-body effects, GSIS was also compromised by P14 in isolated *6F; Pdx1^Cre^* islets (Figure 1E).

**Figure 1:**
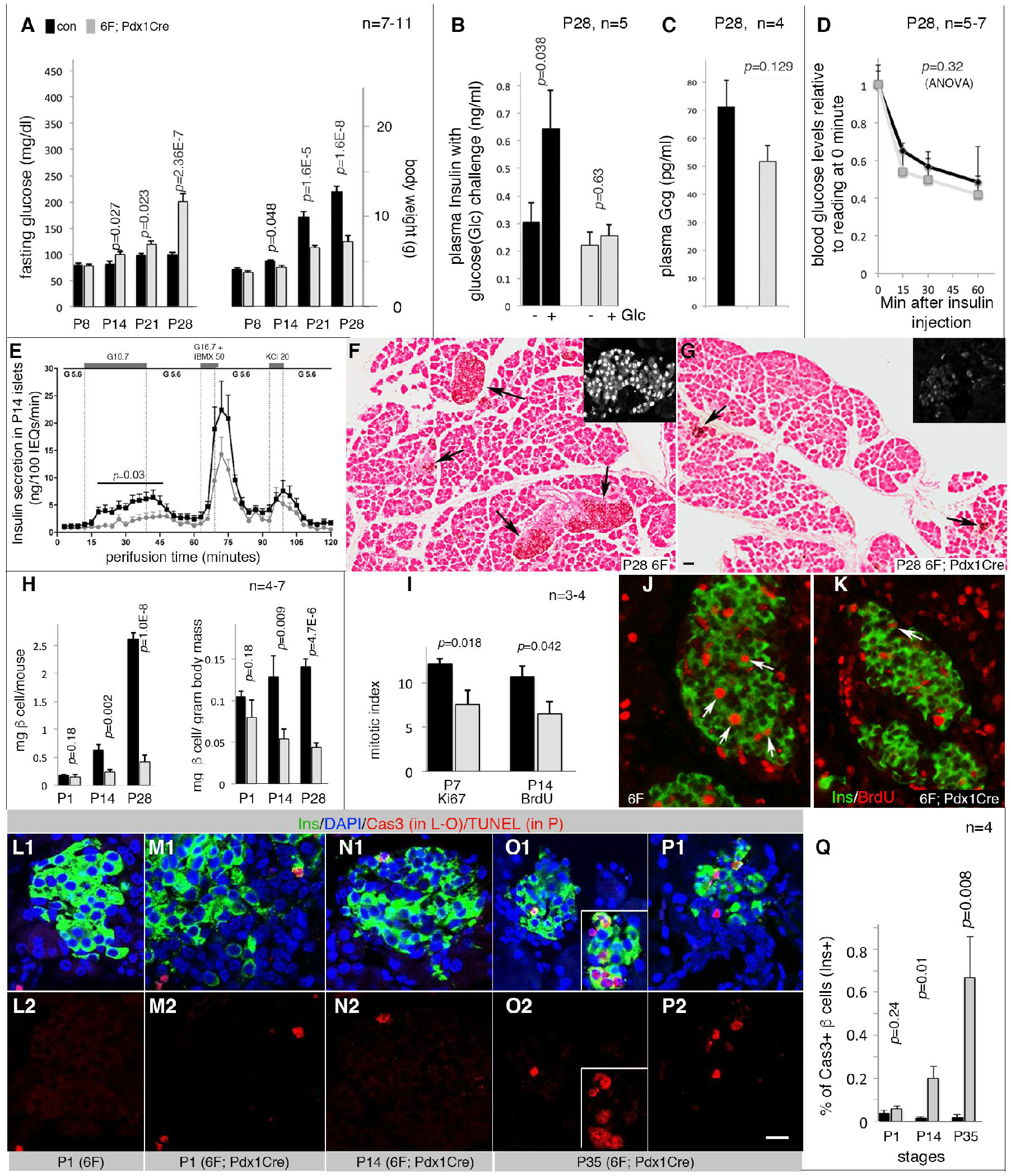
Loss of *Myt* function compromised functional β-cell mass. Mice were derived from inter-cross of *Myt1^F/+^; Myt2^F/+^; Myt3^F/+^; Pdx1^Cre^* breeders or between *6F* and *Myt1^F/F^; Myt2^F/+^; Myt3^F/F^; Pdx1^Cre^* mice. The *p*-values were type 2, 2-tailed t-test except in D and E (ANOVA). Error bars in all quantification panels, SEM. “n”, the number of mice used. Also see Table S1 for data with sexes noted. For panels A-E, H, I, and Q, black bars or lines represent control data. Grey bars or lines represent mutant data. (A) Glycemic phenotypes (left) and body weights (right). The controls include wild-type, *Pdx1^Cre^*, and *6F* mice (see Table S1 for annotation). (B) Plasma insulin levels before and 30 minutes after glucose challenge. (C) Plasma glucagon (Gcg) levels in mice fasted overnight. (D) Insulin tolerance tests. Presented are the blood glucose levels relative to that before insulin injection. *P,* one-way ANOVA. (E) Perifusion-based insulin secretion assays. The track included sequential inductions by: G5.6 (5.6 mM glucose), G16.7, G5.6, G16.7 + 50 μM IBMX (a cyclase inhibitor to increase intracellular cAMP levels), G5.6+KCl (20mM), and ended with G5.6. The *p* value between G16.7-induced secretions is from one-way ANOVA. (F, G) Insulin detection with horse-radish peroxidase (HRP)-staining to visualize Insulin+ cells (brown, arrows). Scale bar = 50 μm. Insets in A and B, examples of islet stained for Pdx1, showing islet morphology. (H) β-cell mass, mg β cells per mouse or mg β cells per gram of body weight. (I-K) Mitotic indices (I) assayed with Ki67 expression (P7, not shown) or BrdU incorporation (P14, J, K. 2-day BrdU feeding) in β cells. Arrows, examples of mitotic cells. (L-Q) β cells with activated Caspase 3 (Cas3, L-O) or TUNEL signals (P). The two rows are merged (top) and single (bottom) channels. Insets in O2 showed a cluster of Cas3^+^ β cells. In panel P2, scale bar = 20 μm, applicable to J-P.

Although *6F; Pdx1^Cre^* mutant islets appeared largely normal in newborn mice (Liu et al., 2019), they became smaller and displayed a much more aspherical morphology at later stages in relation to controls (Figure 1F, G). A reduction in β-cell mass was observed within two weeks of birth (Figure 1H), which accompanied both decreased proliferation (Figure 1I-K) and increased apoptotic cell death (Figure 1L-Q). These findings suggest that the Myt TFs are required for postnatal β-cell function, proliferation, and survival.

### Myt factors are necessary for retaining hallmark features of mature β cells

Molecular defects in perinatal *6F; Pdx1^Cre^* β cells were initially screened via immunofluorescence for various protein products important in their embryonic development and postnatal function, such as the Glut2 glucose transporter and MafA, MafB, Pdx1, and Nkx6.1 TFs (Artner et al., 2006; Fujitani et al., 2006; Matsuoka et al., 2004; Schaffer et al., 2013; Thorens, 2015). At P1, no change was observed in any of these β-cells markers. The levels of Glut2, MafA, and Pdx1 were reduced at the P7 and P28 time-points (Figure 2, columns 1-3). In contrast, Nkx6.1 protein levels were unchanged at all time points (Figure 2, Column 4) while MafB was elevated in β cells after P7 (Figure 2, column 5). We found that the reduction in MafA and Pdx1 TFs in *6F; Pdx1^Cre^* mice was not seen at the mRNA level (Figure S2A), suggesting that the Myt TFs do not transcriptionally control these key β-cell TF genes, and implying a strong post-transcriptional regulatory effect.

**Figure 2.**
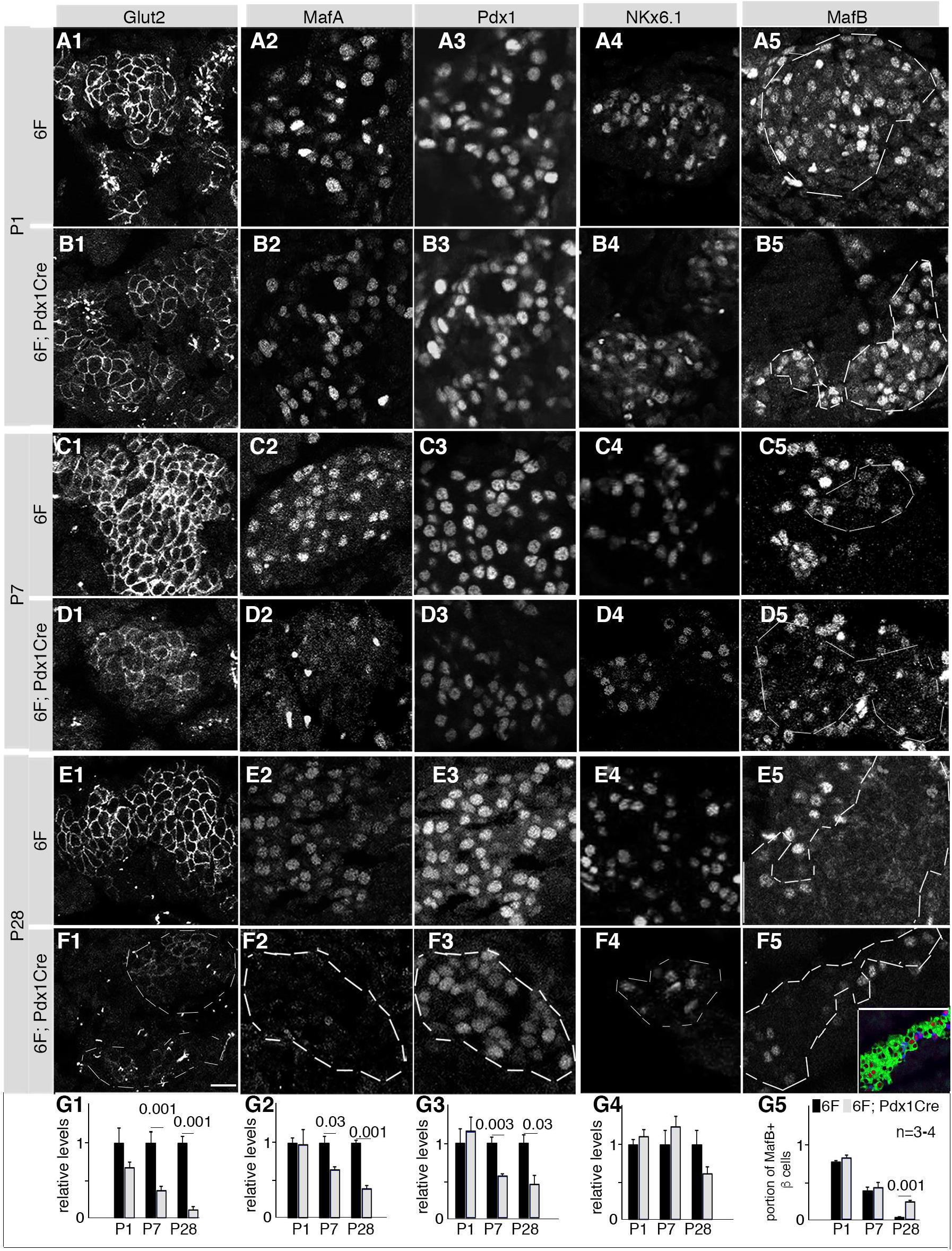
The Myt TFs are required for sustained expression of several β-cell markers. Mice were derived from crosses between *6F* and *Myt1^F/F^; Myt2^F/+^; Myt3^F/F^; Pdx1^Cre^* mice. See Figure S2 for more related expression data. (A-F) Images of each assayed protein. Co-expression of insulin and/or Pdx1 staining was used to locate β cells but not shown [see inset in F5 (green: insulin, red: MafB) for an example]. Dotted-lines in some panels circled β-cell areas (column 5) or islets (F1-F4). Scale bar in F1=20 μm, equal in all panels. (G1-G5) Quantification of relative immunofluorescence of corresponding markers (in each column) at different stage, assayed with Image J. The error bars represent SEM. Three or four mice (“n”) were used for each assay. P values (t-test) smaller than 0.05 were labeled.

### Embryonic inactivation of Myt TFs potentiates stress-gene expression in P1 β cells without activating early-stage stress sensors

To potentially identify direct targets of the Myt-TFs, we next searched for transcriptomic differences between *6F; Pdx1^Cre^* and control β cells. RNAseq analysis of P1 flow-sorted β cells revealed 1,432 down- and 1,432 up-regulated genes (including pseudogenes, Table S2), classifiable into several distinct regulatory pathways by Gene-Set-Enrichment-Analysis (Figure S2B, *p*<0.02, adjusted) (Subramanian et al., 2005). Subsequent analysis focused on stress-response genes in the “*protein processing in ER*” category since these are associated with post-transcriptional control (Pan, 2013) and were amongst the most up-regulated in *6F; Pdx1^Cre^* β cells [see *Hspa1a, Hspa1b, HspH1,* and *Hspb1* expression in Table S2]. Importantly, mRNA expression of 12 of 13 up-regulated stress genes was verified in P1 and P14 *6F; Pdx1^Cre^* islets that had not undergone flow sorting (Figure 3A), eliminating the possibility that sorting stresses – such as the high level of the MIP-driven β-cell–specific eGFP marker and the processing by tissue dissociation and flow sorting – influenced their expression state.

**Figure 3.**
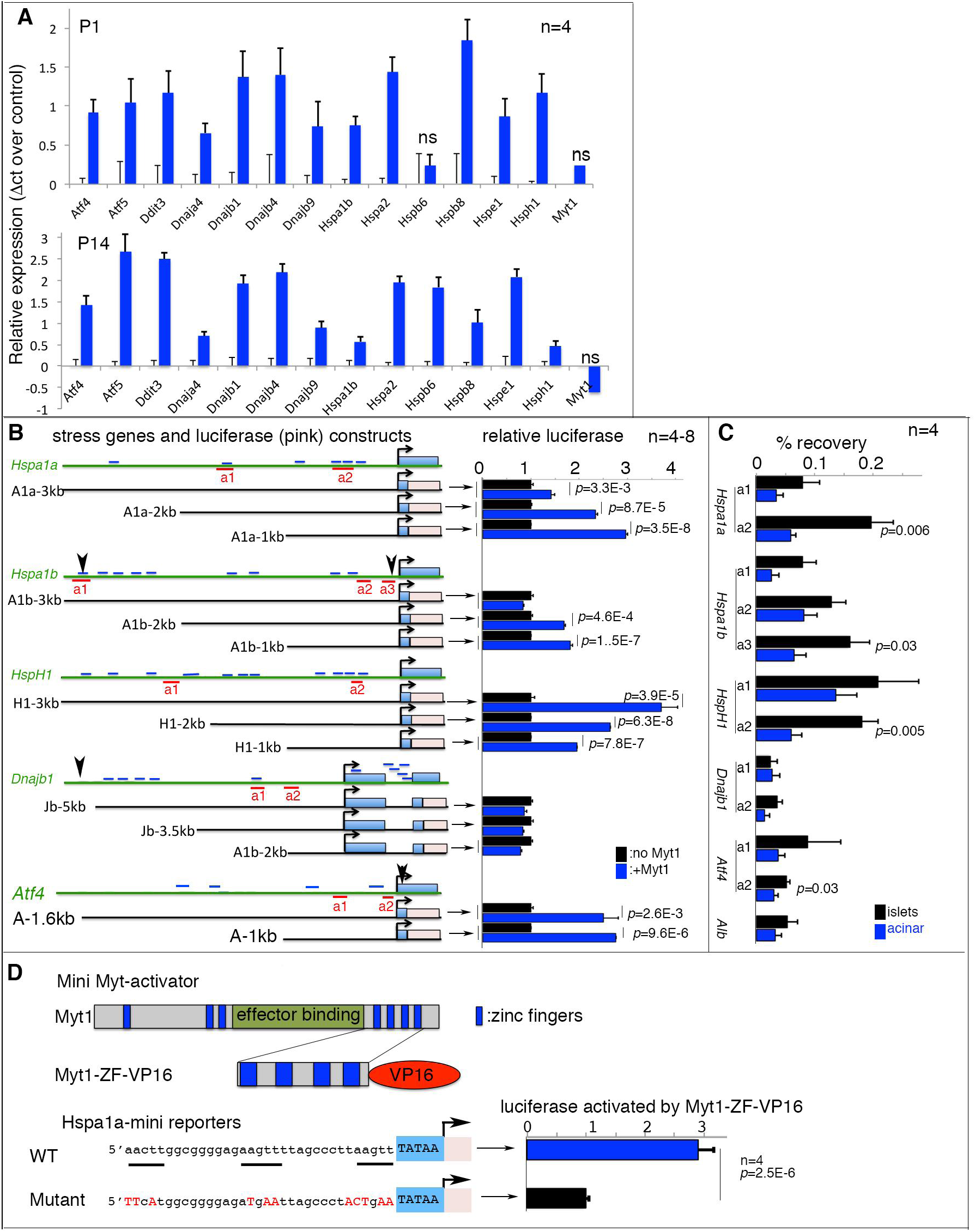
Myt1 binds putative enhancers of several stress-response genes. See Figure S2 and S3 for more related data. Mice were derived from crosses between *6F* and *Myt1^F/F^; Myt2^F/+^; Myt3^F/F^; Pdx1^Cre^* animals. Error bars in all quantification panels, SEM. (A) Real-time reverse transcription-PCR assays of several stress-gene mRNAs in *6F* control and *6F*; *Pdx1^Cre^* islets. “n”, number of mice assayed. All the expression changes except the three marked with “ns” are significant, with *p*<0.05 via T-test (type 2, 2 tails). (B) Reporter assays of stress gene control elements. Diagrams on the left showed the 5’ distal regions of each gene assayed. The approximate length (kilo-base, kb) of used regulatory region was marked. Dark green lines, genomic region of genes. Light-blue rectangles, the stress-gene exons. Pink rectangles, firefly luciferase cDNA. Short blue lines, the AAGTT motifs. Red underlines, the amplicons used for ChIP-PCR in panel C. Arrowheads (*Hspa1b, Dnajb1,* and *Atf4*), DNA elements reported to bind with Myt1 and /or Myt2 in heterologous cells. The right side bars in panel C are relative reporter activities assayed in HEK293T cells. “*”, *p*<0.02 (t-test, type 2, 2-tailed test. 4-8 independent assays). Error bars are SEM. (C) ChIP-PCR assays of Myt1 enrichment on putative stress-gene enhancers. The locations of the PCR amplicons of each gene were marked in panel B. “n” indicates number of immunoprecipitations. The *p*-values (type 2, 2-tailed t-test) smaller than 0.05 were marked. Acinar cells that do not express Myt1 were used as negative controls. (D) Reporter assays using a putative Myt1-binding site on *Hspa1a* enhancers identified in ChIP assays (a2 in panel B). The motif, with wild-type and mutated sequences, was fused to a minimal CMV TATA box to produce reporters. The activator was a fusion protein between the C-terminal 4 zinc-fingers of Myt1 and a VP-16 activation domain. “n”, the number of luciferases assays. “*p*” value is from type 2, 2-tailed t-test.

Despite the up-regulated stress-gene expression in P1 *6F; Pdx1^Cre^* β cells, we detected no stress-associated ER lumen dilation using transmission electron microscopy (Figure S2C), or any general reduction in PERK-eIF2α-regulated protein translation in these P1 islets, although a slight translational reduction was detected at P14 (Figure S2D). In addition, Ire1α-mediated *Xbp1* mRNA splicing was not increased in P1 *6F; Pdx1^Cre^* islets (Figure S2E). Likewise, stress-induced nuclear translocation of Foxo1 was not obvious until P14, when high blood-glucose levels were observed (Figure S2F). Lastly, there was no change in expression of early-acting stress-pathway sensor-effectors that are classical Atf6 targets such as *Xbp1, Hspa5,* or *Hsp90b1* (Figure S2G), nor HSF targets such as the *Cct*s (Figure S2G). Overall, we inferred that the substantial up-regulation in late-acting stress-response gene expression in *6F; Pdx1^Cre^* islets was not mediated by the early-acting UPR sensor-effectors, and consequently postulated that the Myt TFs normally work to directly repress transcription of these late-stage stress-pathway genes. Supporting this hypothesis, *in silico* analysis and data mining of published ChIP-seq data derived from fibroblasts and neuronal progenitors (Bellefroid et al., 1996; Mall et al., 2017; Vasconcelos et al., 2016) have identified several putative Myt TF binding sites (proposed AAGTT consensus, Bellefroid et al., 1996) within 5’ *cis*-regulatory regions of *Atf4*, *Hspa1a*, *Hspa1b, HspH1,* and *Dnajb1* (Figure 3B), although their significance to expression of these stress-pathway genes was not reported.

### Myt1 binds to *cis*-regulatory sequences within stress-response mediators

To determine if the Myt TFs can exert regulatory control via the 5’-flanking control regions of *Atf4, Dnajb1, Hspa1a, Hspa1b,* and *HspH1* genes proposed above, luciferase-reporter expression constructs driven by relatively short sequences spanning these elements were analyzed in co-transfection assays with Myt1 in HEK293T cells (Figure 3B). Myt1 significantly activated expression of the *Hspa1a, Hspa1b, HspH1,* and *Atf4* reporters, but not *Dnajb1*. The more modest effect on *Dnajb1*-driven transcription could reflect the absence in our construct of an additional Myt-binding site detected at roughly -6 kbp in fibroblasts and neuronal cells (Mall et al., 2017)(Vasconcelos et al., 2016), because 5’-flanking control regions containing this putative binding site were for unknown reasons unclonable. Note that while we were probing the mechanisms for the transcriptionally repressive effects of Myt TFs on stress-pathway genes that we had detected in pancreatic β cells, these out-of-context assays in HEK293T cells showed activation not repression, which we expected based on the similar “reversals” that occurred when testing the AAGTT motifs in artificial *in vitro* reporter-transfection assays (Bellefroid et al., 1996; Manukyan et al., 2018; Yee and Yu, 1998).

ChIP-PCR assays using islets were used to test directly for Myt1 binding within *Hspa1a*, *Hspa1b*, *Hsph1, Dnajb1*, and *Atf4*. Only Myt1 binding was analyzed for efficacy, as its zinc-finger DNA-binding region shares over 80% identity with that of Myt2/Myt3, and Myt1 antibodies are of high quality (Figure S3A). Acinar-cell chromatin, which does not express detectable Myt1, was the negative control (Figure S3B). Significant binding was observed within several Myt1-element– containing regions of *Hspa1a*, *Hspa1b*, *Hsph1,* and *Atf4* (Figure 3C).

Myt1 binding was co-verified using a reporter driven by the Myt1 binding elements within the -435 to -402 bp region in *Hspa1a*. Myt1 responsiveness was analyzed using wild-type and mutated Myt-binding site constructs in cotransfection with a Myt1 DNA binding domain fused to the VP16 transactivation domain (i.e. Myt1ZF-VP16). Activation by Myt1ZF-VP16 in HEK293T cells was dependent on the presence of the AAGTT motifs (Figure 3D). Our data in pancreatic β cells, combined with the ChIP-seq data from neuronal progenitors and fibroblasts (Mall et al., 2017)(Vasconcelos et al., 2016), leads to our supposition that Myt TF–based regulation of stress-pathway genes is functionally relevant in a variety of cell types.

### Over-expressing *Hsps* or *Atf4* compromises postnatal islet β-cell proliferation, survival, and function

A tetracycline-inducible transgene was used to over-express HspH1, Hspa1b, Dnajb1, and eGFP [(termed *TetO^3H^*, Figure S4A); eGFP serves to identify the overexpressing cells]. The transgenic mouse driver to achieve HSP overexpression (Hsp-OE) was *Rip^rTTA^*, which produces the reverse tetracycline-controlled transactivator specifically in β cells (Nir et al., 2007). Continuous doxycycline (Dox) administration in the drinking water from E16.5 resulted in ∼6-fold Hsp-OE in P2 *Rip^rTTA^*; *TetO^3H^* islets (Figure S4B). As expected from reports (Brennand et al., 2007; Cai et al., 2012; Dadi et al., 2014), *Rip^rTTA^* only activated Hsp-OE in a portion of the β-cell pool, producing mosaic islets with intermixed overexpressing (eGFP^+^) and normal (eGFP^−^) cells (Figures 4A and S4C).

**Figure 4.**
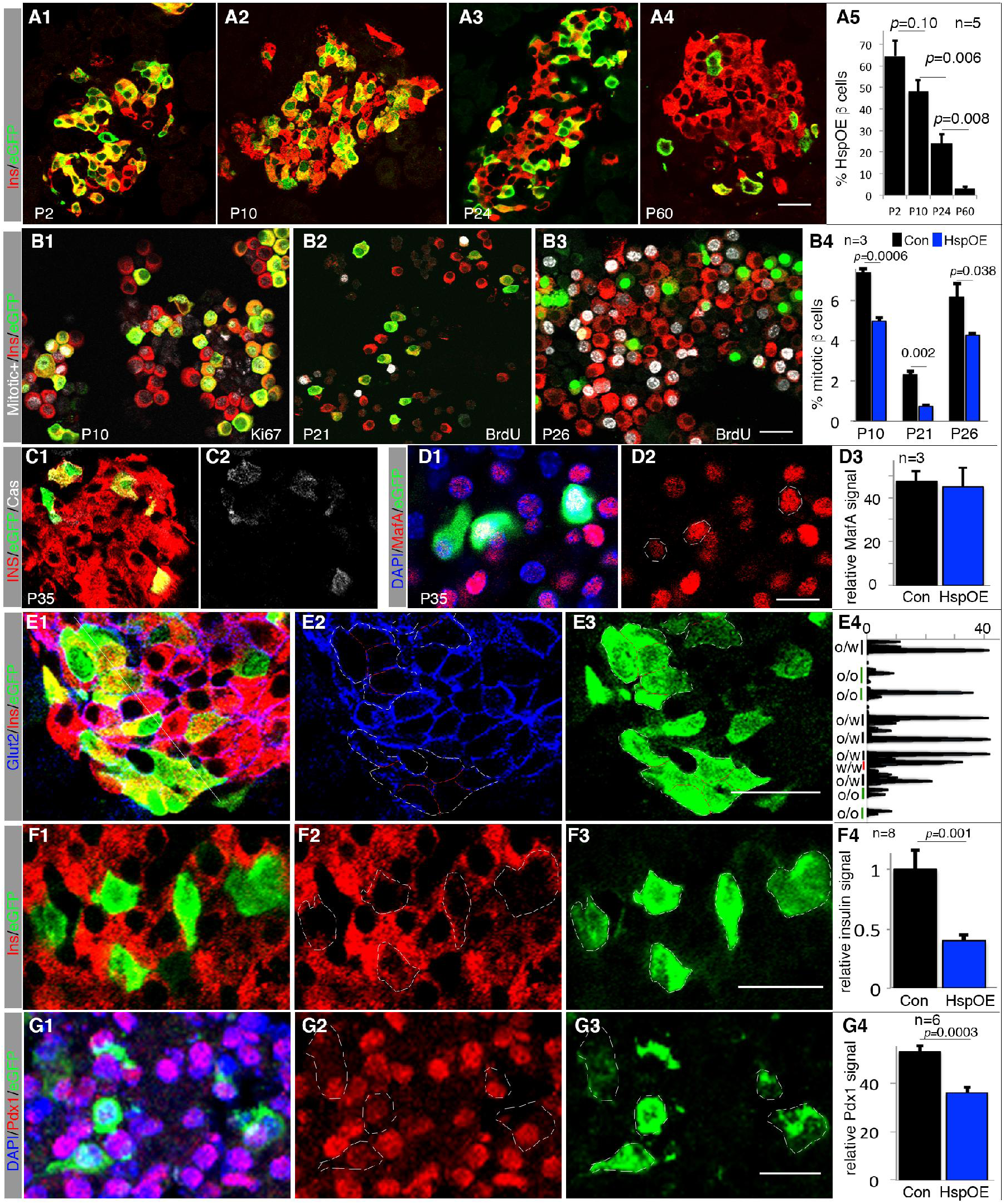
*Hsp* over-expression (OE) compromised β-cell proliferation, identity, and survival. See Figure S4 for other characterization of *Hsp-OE* mice. Error bars in all quantification panels, SEM. In all quantification panels, “n”, number of pancreata examined. (A) The portion of Hsp-OE β cells at several postnatal stages, monitored via the eGFP expression. The *p*-values (type 2, 2-tailed t-test) were marked. (B) Mitotic activity of β cells with (eGFP+ cells) or without Hsp OE (eGFP-cells). P10 (B1) cells were assayed via Ki67 expression. P21 (B2) and P26 (B3) cells via BrdU incorporation after BrdU feeding from water supply between P19-P21 or P21-P26, respectively. Scale bar=20 μm. The *p*-values (type 2, 2-tailed t-test) were marked. (C-G) Expression of β-cell markers in P35 islets with Hsp OE. Merged and single channels were shown, with Image-J-aided quantification. E4 is a line scan (yellow thin line in E1, from top-left to bottom-right), showing the Glut2 intensity along the borders of over-expressing (O) and non-over-expressing (W) cells. The P-values (type 2, 2-tailed t-test) smaller than 0.05 were marked on top of each assay. The circles in C2, D2, E2, and E3 mark the outline of whole β cells. Scale bars = 20 μm.

The proportion of eGFP^+^ β cells in Hsp-OE mice fell from ∼64% at P2 to 1-3% at P60 under continuous Dox-induced Hsp OE (Figure 4A). There was no transgene silencing (Figure S4D, E), but lowered proliferation rate in pre- and post-weaning Hsp-OE cells (Figure 4B). There was also increased apoptotic cell death by P35 (Figure 4C), although not at P10 (Figure S4F), indicated by detection of activated Caspase 3 in most Hsp-OE but not non-OE cells. While the MafA and MafB protein levels were not affected by Hsp-OE at several stages examined (Figure 4D, S4G, H), the levels of Glut2, insulin, and Pdx1 were reduced in P35 (Figure 4E-G) but not P10 Hsp-OE β cells (Figure S4F-I). These results suggest that Hsp-OE recapitulates a portion of the *6F; Pdx1Cre* phenotypes in proliferation, cell survival, and expression of some but not all β-cell factors.

To determine if Atf4 upregulation is responsible for the other portion of the *6F; Pdx1Cre* phenotypes, ATF4-OE was activated by crossing *Ins1^Cre^* with *Rosa26-ATF4^LoxTG^* mice. This resulted in ∼3-fold ATF4-OE in P1 islets (Figure S5A-B). While mice with ATF4-OE displayed only slightly higher blood-glucose levels compared with controls at P8, most developed diabetes by five weeks of age (Figure S5C). These defects were preceded by an increased α- to β-cell ratio and decreased Glut2, insulin, MafA, and Pdx1 production in β cells (Figure 5A-E and data not shown). The relative change in α to β cells coincided with increased α and decreased β-cell proliferation (Figure 5F1-F3), without obvious β-cell apoptosis in 1-month or 3-month old ATF4-OE islets (Figure S5D and data not shown). Additionally, ATF4-OE islets secreted an abnormally high amount of insulin at non-stimulating (basal) glucose concentrations, although substantial glucose-responsiveness was maintained (Figure 5G). Our observation is consistent with *Atf4* inactivation potentiating GSIS (Yoshizawa et al., 2009).

**Figure 5.**
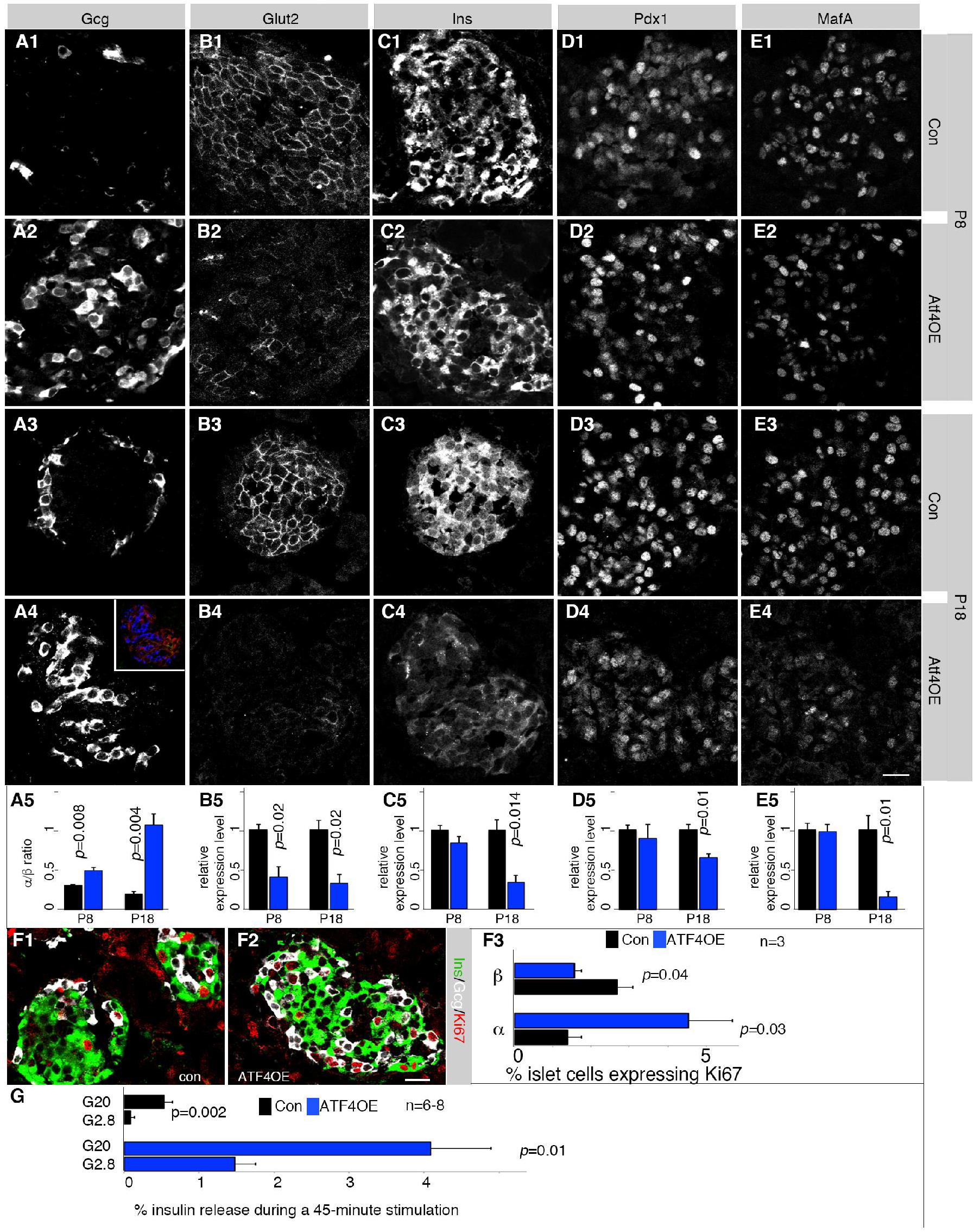
*ATF4* OE compromised β-cell gene expression and insulin secretion. *ATF4* OE was initiated in β cells using the *Ins1^Cre^* and *Rosa26-ATF4^LoxTG^* mouse lines. See Figure S5 for crossing and other characterization of *HSP-OE* mice. Error bars, SEM. (A-E) Detection of several β-cell markers. Controls are *Rosa26-ATF4^LoxTG^* pancreata. Inset in D1 is an example to show how α (blue) and β (red) cells were identified. In E, three to four (n=3-4) pancreata were quantified. The P-values (type 2, 2-tailed t-test) smaller than 0.05 were marked. Scale bar in D5 = 20 μm, applicable to all panels. (F) Mitotic assays via Ki67 staining. “n”, number of pancreata quantified. The P-values (type 2, 2-tailed t-test) were marked. Scale bar = 20 μm. (G) GSIS assays of P14 control and *ATF4-OE* islets. “n”, the number of secretion assays, from 3 mice (with each has at least two technical duplicates). The P-value is from type 2, 2-tailed t-test.

Also to argue against the overexpression of any protein non-specifically compromising β-cell proliferation and/or gene expression, we over-produced tdTomato (tdT) in pancreatic β cells using *Pdx1^CreER^; Ai9* mice. Tamoxifen administration at E15.5 led to activation of *tdT* expression in ∼50% of pancreatic β cells (Liu et al., 2013). Unlike the OE of HspH1, Hspa1b, and DnajB1 or ATF4, tdT had no detectable effect on adult insulin or Pdx1 levels (Figure S5F). Collectively, these results demonstrate that the Myt TFs normally prevent the over-activation of *Atf4*, *HspH1*, *Hspa1b*, and *DnajB1*, which is consequential to postnatal mouse β-cell proliferation, survival, and function.

### Myt3 protein levels are induced in response to metabolic stress in diabetic *db/db* mouse β cells

While inactivation of Myt TFs leads to dysregulated stress pathways, we also tested if there is a reciprocal tuning of Myt TFs levels in response to physiological stresses that are connected to stress responses, such as increased insulin secretion in obese diabetic mice. Comparing islets from 3-month old *db*/+ control (euglycemic) and *db/db* diabetic mice, we found no change in Myt1 or Myt2, but Myt3 protein levels were significantly up-regulated in *db/db* β cells (Figure 6A-D). Intriguingly, *Myt1, Myt2,* and *Myt3* transcript levels were not changed (Figure 6E), suggesting that the increased Myt3 protein level was acquired via post-transcriptional mechanisms. Notably, isolated *db/db* islets were glucose responsive despite the overall hyperglycemic state of the mice (Figure 6F). These findings are consistent with a model that stress-responsive Myt3, together with constitutive Myt1 and Myt2, protect β cells under metabolic stress.

**Figure 6.**
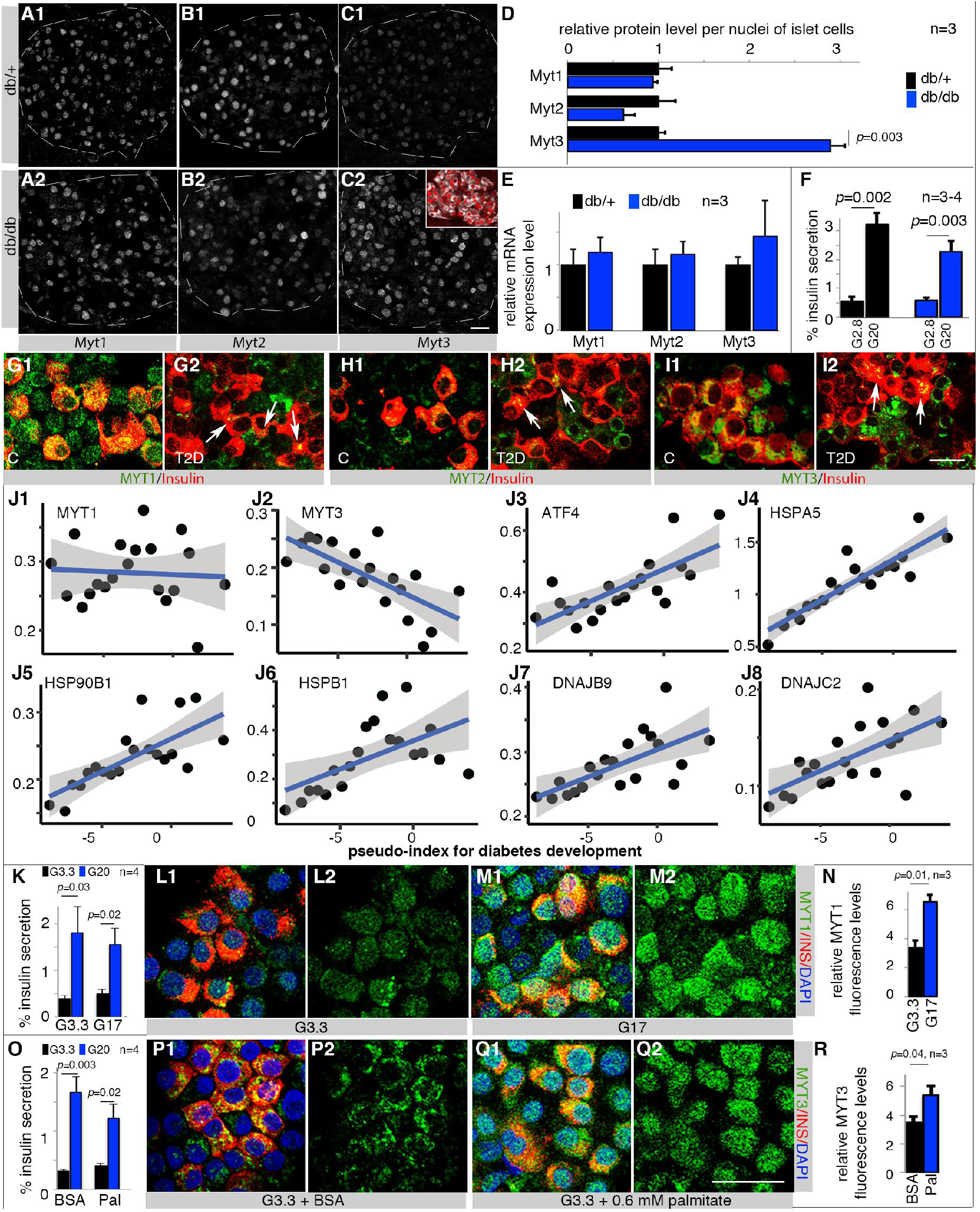
Metabolic stress upregulates Myt (MYT) TF proteins in mouse and human β cells while T2D downregulates *MYT TFs* but upregulates stress-gene transcripts. See Figure S6 and Table S3 for more characterization of human islets. Error bars in all quantification panels, SEM. (A-F) Myt TF expression at protein (A-D) and mRNA (E) levels in β cells/islets of *db/+* and *db/db* mice that maintain GSIS (F). Inset in C2 showed β cells (Insulin+, white) that express Myt3 (red). “n”, number of mice examined. The P-values (type 2, 2-tailed t-test) smaller than 0.05 were marked. Scale bar in C2 = 20 μm, applicable from A-C. (G-I) MYT TF levels in functional (G1, H1, and I1) and non-functional (G2, H2, and I2, arrows point to several MYT+ β cells) human islet β-cells. Representative results from three donors, with merged channels between MYT and Insulin signals shown. Scale bar = 20 μm. (J) RePACT-analysis of *MYT* and stress gene transcription in ∼35,000 β cells of six normal and 3 T2D islet donors. X-axis, pseudotime trajectory of T2D progression (left to right). Y-axis, relative gene expression. Each dot represents a β-cell subpopulation. The blue line represents average gene expression trajectory. The blue shades indicate range of gene expression with 95% confidence levels. (K-N) GSIS [K, under 3.3 mM (G3.3) and 20 mM (G20) glucose] and MYT1 production of human β cells after 40-hour treatment of 17 mM (G17) glucose. “n” in panel K, number of GSIS assays. “n” in N, batches of islets used for MYT1 quantification. A merged MYT1/Insulin and a single MYT1 channel were shown. The marked p-values were type 2, 2-tailed t-test. Scale bar = 20 μm. (O-R) Same as K-N except 0.6 mM palmitate (Pal) was used instead of G17. (R)

### Downregulated MYT-TF levels are associated with stress-gene upregulation in human T2D islet β cells

The MYT1, 2, and 3 proteins were immunodetected in most functional human islet β cells (Figure 6G-I). While MYT1 mostly appeared nuclear (Figure 6G1), MYT2 and MYT3 were cytoplasmic (Figure 6H1, I1). These factors were also substantially downregulated in most β cells of T2D human donors (Figures 6G2, H2, I2), which were dysfunctional by whole-islet GSIS analysis (Figure S6A). However, some cytoplasmic MYT TF was still observed in some β cells (Figure 6G2, H2, I2, arrows).

MYT TF mRNA expression was next examined in T2D human β cells. A large challenge for this analysis is the heterogeneous nature of T2D, both between individual patients and between islets from the same patient, previously proposed to account for the poorly overlapping data between various samples (Wang and Kaestner, 2019). To avoid this issue, we utilized our recently devised RePACT algorithm to analyze single-cell expression data, which can reveal progressive movement along a pseudotime-trajectory from functional to dysfunctional β-cell state (Fang et al., 2019). There was a significant anti-correlation (adjusted *p*=0.0007) between the RePACT-inferred T2D-progression process and *MYT3* mRNA levels, but not for MYT1 or MYT2 (Figure J1, J2, and Table S3). A positive correlation was observed between the T2D pseudo-timeline and expression of several *ATFs* (Figure 6J3, and Table S3), *HSPA5* (BIP), *HSP27, 40,* and *90* family genes (Figure 6J4-J8 and Table S3). These results provide further evidence for (at least) MYT3 influencing stress-gene expression in human β cells. They also imply post-transcriptional mechanisms in regulating the MYT TF levels, as seen for Myt3 in *db/db* islets

The reduced MYT TF expression levels in T2D β cells likely result from the long-term effects of hyperglycemia and/or hyperlipidemia (Swisa et al., 2017). We therefore investigated the acute effects of both stressors. Treating human islets for 40 hours at a supra-physiological 16.7 mM glucose concentration had little effect on GSIS (Figure 6K), but significantly and selectively up-regulated the MYT1 protein level in islet β cells (Figure 6L-N, S6B, C). In contrast, lipotoxic 0.6 mM palmitate treatment, which compromised but did not eliminate GSIS (Figure 6O), rather specifically increased the level and nuclear localization of MYT3 [Figure 6P-R; compared to MYT1 and MYT2 (Figure S6D, E)]. Again, *MYT1* or *MYT3* transcript levels were unchanged by either treatment (Figure S6F), which further supports the significance of post-transcriptional mechanisms in regulating MYT TF protein levels. Thus, pathophysiological effectors of T2D β cell dysfunction can influence MYT TF protein levels in human islets – most predominantly through post-transcriptional effects on MYT3.

### MYT TFs are required for human β-cell survival

To provide further support for MYT TF action in stress-pathway gene regulation in human β cells, siRNA-based *MYT* knockdown was performed in the human EndoC-βH1 cell line, which displays functional properties similar to primary human islet β cells (Tsonkova et al., 2018). MYT1, 2, and 3 were all detected and MYT siRNA treatment effectively reduced their protein and mRNA levels in a majority of EndoC-βH1 cells (Figure 7). These conditions led to elevated *ATF4*, *DNAJB1*, *HSPA1*, and *HSPH1* expression (Figure 7D) and increased β-cell death (Figure 7E), strongly supporting the integral and conserved role for the MYT TFs in stress-gene regulation in islet β cells.

**Figure 7.**
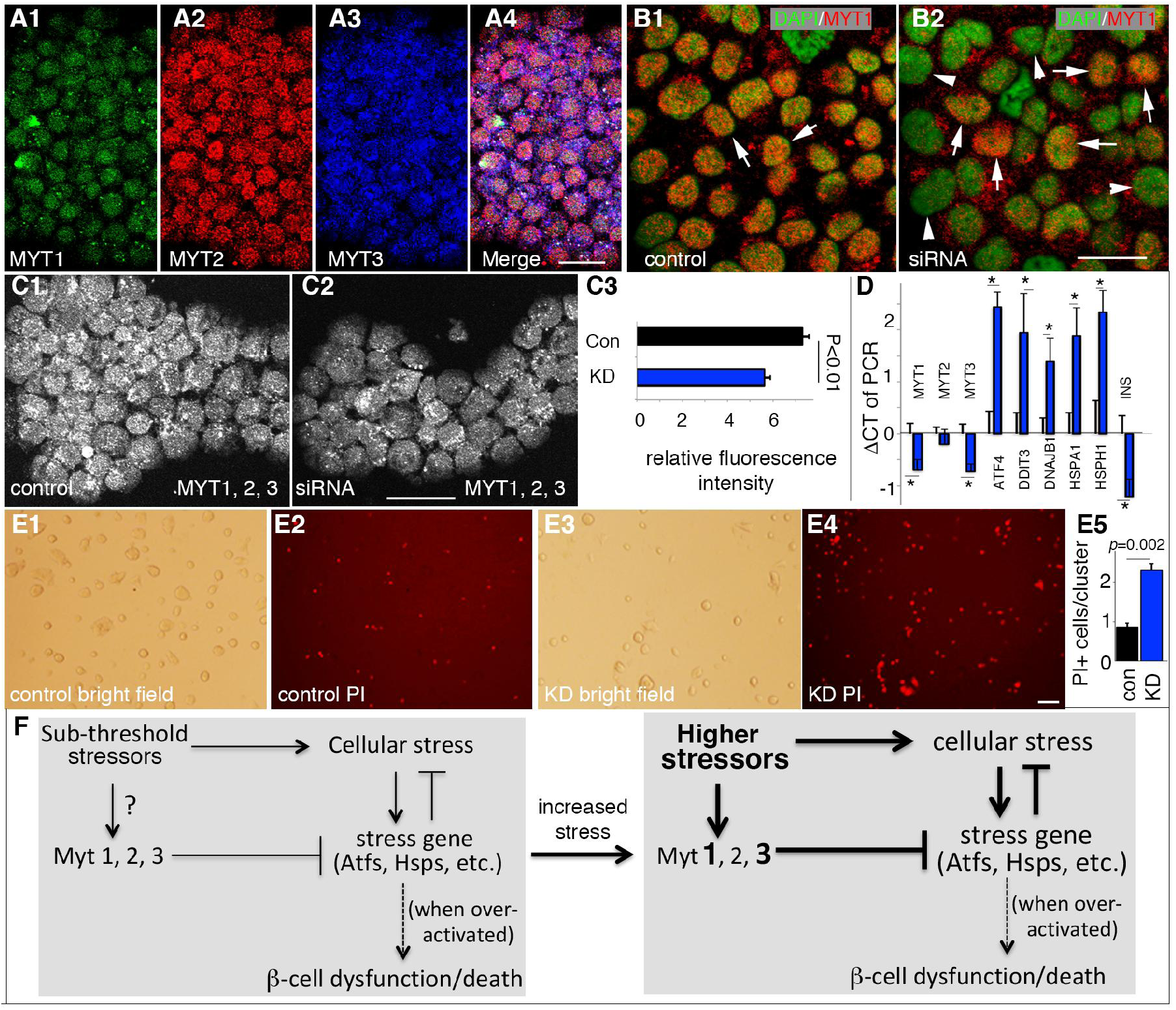
MYT TFs is required for human β-cell line survival. All assays were done three days after cell re-plating. Error bars in all quantification panels, SEM. (A) MYT TF detection in EndoC-βH1 cells. Individual and merged channels were all presented. Scale bar = 20 μm. (B) Representative images of siRNA-based *MYT1* KD three days after siRNA transfection. Arrows, examples of MYT1^+^ nuclei. Arrowheads in B2, examples of cells with low MYT1 signal. Scale bar = 20 μm. (C) Staining and quantification of MYT TFs in viable EndoC-βH1 cells after *MYT* KD. MYT1, 2, 3 signals were combined into one single channel (C1, C2) and quantified with Image J (C3). The *p*-value was from type 2, 2-tailed t-test. Scale bar = 20 μm. (D) Effects of *MYT* KD on the transcription of several stress genes in EndoC-βH1 cells. “*”, *p*<0.05, from type 2, 2-tailed t-test. (E) Propidium Iodide (PI) assays of cell death three days after siRNA KD. Bright field and PI fluorescence were shown (E1 and E2, a field transfected with control RNA. E3 and E4, a field transfected with MYT siRNA). The quantification was by counting the number of PI+ cells per cell cluster. Scale bar = 100 μm. The *p*-value was from type 2, 2-tailed t-test. (F) A model depicting Myt functions in β-cell stress response. Under normal conditions (left box), sub-threshold stressors induce low level of cellular stress and stress-gene expression, which reduce cellular stress. The Myt TFs, may or may not rely on stress for expression, ensure low stress-gene expression to prevent β-cell dysfunction and death. Higher stressors (right box) induce higher cellular stress and stress-gene expression for protein homeostasis while also induced Myt1/Myt3, which again prevent late-stage stress-gene over-activation and β-cell dysfunction and/or death.

## Discussion

Stress-survival response pathway activation needs to sufficient to allow proteomic homeostasis (misfolded protein clearance, chaperone activity, etc.) but not so high as to induce catastrophic cellular dysfunction and death. Such tuning is particularly important for islet β cells which cannot be readily regenerated but produce stressors as part of their normal physiological function (Cao and Kaufman, 2014; Farley and Watkins, 2018; Fulda et al., 2010). Several molecular pathways by which cells activate stress-pathway gene expression have been established (Cao and Kaufman, 2014; Ma, 2013; Papa, 2012). Yet how cells maintain the appropriate amplitude to assure physiological, and not pathophysiological, expression is undetermined. We show here that the Myt TF family is required in controlling the response of β cells to stress (Figure 7F). Given that ChIP analysis revealed Myt TF sites within lkely transcriptional control sequences of the *Hspa1a*, *Hspa1b*, *Hsph1,* and *Atf4* genes in islet β cells, neuronal progenitors (Vasconcelos et al., 2016)(Mall et al., 2017) and fibroblasts (Vasconcelos et al., 2016), it is possible that Myt TFs are involved in a general protective mechanism that allows stress-prone cells to function and survive.

### Myt TFs regulate several signaling pathways

We have uncovered several cellular processes that the MYT TFs control in β cells. Our earlier report of their inhibition of precocious *Syt4* expression and, consequently, regulation of Ca^2+^ signaling and insulin secretion (Huang et al., 2018), adds to the present connection to the stress-response of postnatal β cells. Additionally, we showed that loss-of *Myt* function deregulates over a dozen other genetic pathways (e.g., PI3K-AKT, Rap1, and Ras signaling; Figure S2B) that have well-established roles in β-cell proliferation and function (Elghazi et al., 2007; Font de Mora et al., 2003; Kelly et al., 2010). Given that manipulating *Syt4* or stress-gene expression levels only partially recapitulates the overall phenotype arising from combined *Myt1/Myt2/Myt3* loss-of function, we propose that the Myt family could be considered as a functional nexus that interconnects these intracellular responses to developmental and physiological signals.

### Bidirectional connections between Myt TFs and stress-response pathways

The absence of Myt TF in *6F; Pdx1^Cre^* P1 islet β cells resulted in precocious over-expression of late-stage stress-pathway genes, but they lacked concurrent signs of cellular stress response such as ER dilation or activation of some signature UPR and OSR genes. The conclusion is that the Myt TFs do not regulate the early-stage stress sensor/effectors (e.g., IRE1 and PERK), which can be activated for proteomic homeostasis in the presence of upregulated Myt TFs. Instead, these TFs mainly prevent the overactivation of the late-stage effectors (e.g., Atf4) that are more likely to trigger stress-incuced cell death/dysfunction (Hetz and Papa, 2018). Our ChIP and Myt-motif–driven transfection assays together strongly suggest that regulation is mediated by direct binding to 5’-flanking *cis*-regulatory sequences for at least *Hspa1a, Hspa1b, HspH1,* and *Atf4*, in agreement with results for Myt1 or Myt2 in fibroblasts and neuronal progenitors (Mall et al., 2017; Vasconcelos et al., 2016).

Our studies have revealed a complex cross-regulation between Myt TF and stress. In both murine and human β cells, stress conditions that did not eliminate GSIS caused upregulation of MYT TF protein levels via post-transcriptional mechanisms. Specifically, this was observed in the context of Myt3 protein upregulation in *db/db* obese mice, and MYT1 and MYT3 protein production and/or nuclear localization enhancement in human β cells under short-term high glucose or palmitate treatment, respectively. Intriguingly, dysfunctional β cells from T2D human donors had significantly reduced *MYT3* and upregulated stress-gene transcription. This latter point, supported by our RePACT-based single-cell RNAseq, is consistent with results from conventional gene expression analyses, which showed down-regulation of MYT3 (Axelsson et al., 2017) and up-regulation of *ATF1, ATF3, Hsp40, 70, 90, and 105* (Bugliani et al., 2013; Gunton et al., 2005; Hartman et al., 2004; Marselli et al., 2010; Segerstolpe et al., 2016).

Collectively, our findings support the following model (Figure 7F). In β cells not suffering abnormally high-stress conditions, the Myt TFs ensure relatively low basal stress-gene expression. During metabolic stress, early-stage stress sensors/effectors (IRE1, PERK, Atf6, Nrf2, and etc.) were activated to maintain proteomic homeostasis (Cao and Kaufman, 2014; Hetz and Papa, 2018). The stressors also up-regulate Myt1/Myt3 protein levels to prevent over-activating the late-stage stress pathways (e.g., Atf4 and Hsps), which has the potential for inducing catastrophic β cell failure and death. These mechanisms are only partially protective, because sustained stress can nonetheless cause its disablement, indicated by the reduced MYT TFs in T2D β cells. This situation is notably similar to Foxo TFs, which become activated during early β-cell compensation but are later inactivated, for unknown reasons, during β cell failure (Accili et al., 2016). It is plausible that exploring the mechanisms of Myt family regulation, perhaps first focused on the transcriptional and post-transcriptional regulation of MYT3, may provide ways of targeting MYT TFs to offset or prevent β cell failure and T2D.

### Myt TFs regulate a subset of stress-response genes

All stress-regulated genes are not upregulated in *Myt-*deficient β cells. For instance, *Hspa5, superoxide dismutase* (*Sod*) and *catalase* (*Cat*) were unaltered in *6F; Pdx1^Cre^* compared to control β cells. This profile distinguishes the Myt TFs from other stress-activated TFs such as Foxo1 and Nrf2. Foxo1 potentiates *MafA*, *Pdx1*, ROS scavenging enzyme gene (i.e. *Gpx*, *Sod*, *Cat)* transcription to improve glucose sensing, insulin production, cell proliferation, and ROS removal (Kitamura et al., 2005; Talchai et al., 2012; Zhang et al., 2016). Nrf2 activation elevates the expression of antioxidant proteins for ROS degradation (Ma, 2013). We propose that the non-completely overlapping responses of Myt, Foxo, and Nrf2 TFs to cell stressors collectively provides for multimodal selectivity amongst the various kinds of cellular stress-survival machinery.

### Myt TFs may have both repressor and activator activities

While Myt TFs were believed to predominantly repress gene transcription *in vivo* (Mall et al., 2017; Manukyan et al., 2018; Vasconcelos et al., 2016), ChIP-seq studies identified Myt1 or Myt2 motifs that were associated with activation by *in vitro* assays (Mall et al., 2017; Vasconcelos et al., 2016), suggesting a strong context-dependent ability for Myt TFs to act as activators or repressors. We found equal numbers of up- and down-regulated genes in *6F; Pdx1^Cre^* β cells from newborn mice (Table S2). Intriguingly, many up-regulated genes fell into the stress-response category, whereas down-regulated genes were generally associated with promotion of β-cell proliferation and function (Figure S2B). Thus, we speculate that Myt TFs can both directly activate genes required for β-cell compensation and repress those mediating cell failure. This proposal should be testable by orthogonal ChIPseq and RNAseq analysis in primary β cells.

Precisely how Myt TFs regulate transcription is unknown, although the Sin3 and Lsd1 co-regulators have been inferred as functional partners (Mall et al., 2017; Vasconcelos et al., 2016). The co-repressor Sin3, which has Sin3A and Sin3B paralogs, can recruit histone deacetylases (Grzenda et al., 2009) and was reported to interact with Myt TFs in neuronal and pancreatic β cells (Romm et al., 2005; Scoville et al., 2015). Another repressor, Lsd1, or lysine-specific histone demethylase 1, was detected in a Myt1-containing transcriptional complex in neuronal cells (Yokoyama et al., 2014). Because Myt TFs may recruit Sin3 and/or Lsd1 to effect transcriptional repression (Vasconcelos et al., 2016; Yokoyama et al., 2014), direct experimentation, presumably starting with ChIP-type approaches, may distinguish the respective roles of Sin3 and Lsd1 and if they operate on independent or overlapping sets of genes.

In summary, we have uncovered a previously unknown role for the Myt TFs in controlling cellular response to stress in pancreatic islet β cells, but potentially extendable to other cell types. Our data further suggest that the degree of repression tailors the stress response within a functionally favorable range to allow long-term cellular function and survival.

## Acknowledgement

We thank Chunhua Dai and Alvin C. Powers of Vanderbilt Universtity Medical Center for assistance with the human islet studies. This study is supported by grants from NIDDK (DK065949 for GG and RS) and JDRF (1-2009-371 for GG). Confocal and TEM imaging were performed with VUMC Cell Imaging Shared Resource (supported by NIH grants CA68485, DK20593, DK58404, DK59637 and EY08126). We also thank the Islet Isolation and Procurement Core of Vanderbilt Medical Center for hormone assays (funded by DK20593).

## Author Contributions

G.G., Y.L., R.S, A.N.B, and C.V.E.W. conceptualized the work and designed the experiments. R.H., Y.X., C.H., G.E.E., M.B., and G.G. did mouse characterization, gene expression, imaging, and secretion assays in mice. A. G. and M.B. did islet perifusion. R.H. and G.G did the reporter and ChIP assays in islets. E.W. and R.S. did *MYT* knockdown in human β cells. C.W. and L.Y. analyzed gene expression in human β cells. A.N.B., Y.X., and GG prepared primary human islets. All authors participated in manuscript writing.

## Materials & Correspondence

Correspondence and material request should be addressed to: (615-936-3634).

## STAR methods

### Mice derivation and usage

Mouse usage followed protocols approved by the Vanderbilt University IACUC for GG, in compliance with regulations of AAALAC. All mice were euthanized by isoflurane inhalation, followed by decapitation or cervical dislocation.

Wild-type CD1 (ICR) mice were from Charles River Laboratories. The *C56BL/6J*, *Ai9* [Gt(ROSA)26Sor^tm9(CAG-tdTomato)Hze^], *Rip-rTTA* [Tg(Ins2-rtTA)2Efr/J]*, Rosa26-ATF4^LoxTG^* (B6;129X1-*Gt(ROSA)26Sor^tm2(ATF4)Myz^*/J), and *Ins1^Cre^* [B6(Cg)-*Ins1^tm1.1(cre)Thor^*/J] mice were purchased from the Jackson laboratories. The *Pdx1^Cre^* and *Pdx1^CreER^* mice were described in (Gu et al., 2002). The derivation of *Myt1^F/+^, Myt2^F/+^,* and *Myt3^F/^*^+^ mice were described in (Huang et al., 2018; Wang et al., 2007). The *TetO^3H^* mice were derived by pronuclear injection, with a DNA construct made by ligating a bi-directional Tet-ON-3G promoter (Clontech), the coding sequences of *Hspa1b*, *HspH1*, *Dnajb1*, *eGFP* and SV40 early poly-adenylation signals as indicated in Figure S4A. The entire construct was sequence-verified. Note that H2A peptide-breakers were included so that each mRNA could make two proteins (Figure S4A). Five independent transgenic lines were derived, with founders directly crossing with *Rip^rTTA^* mice to establish stable lines.

For producing the Myt triple mutants, *Myt1^F/F^; Pdx1^Cre^* mice [∼50% CD1 background, based on crossing history (Wang et al., 2007)] were crossed with Myt2^F/+^ mice [∼50% CD1 background]. *Myt1^F/+^; Myt2^F/+^; Pdx1^Cre^* mice were then crossed with *Myt3^F/+^* animals (∼50% CD1 background) to obtain *Myt1^F/+^; Myt2^F/+^; Myt3^F/+^; Pdx1^Cre^* mice, which were out-crossed twice with CD1 mice in order to ensure a mixed CD1 genetic background. The *Myt1^F/+^; Myt2^F/+^; Myt3^F/+^; Pdx1^Cre^* mice were then intercrossed. From ∼20 litters of mice, two wild-type (WT), four *Pdx1^Cre^*, five *Myt1^F/F^; Myt2^F/F^; Myt3^F/F^* (denoted as *6F*), and four *Myt1^F/F^; Myt2^F/F^; Myt3^F/F^; Pdx1^Cre^* (denoted as *6F; Pdx1^Cre^*) mice were obtained. No phenotypic differences were found amongst the WT, *Pdx1^Cre^,* and 6F mice (Figure 1 and Table S1). Thus, most of the studies use *6F* (littermates of *6F; Pdx1^Cre^* mice) as controls, derived from two types of crosses: 1) *6F* X *Myt1^F/+^; Myt2^F/F^; Myt3^F/F^; Pdx1^Cre^*; 2) *6F* X *Myt1^F/F^; Myt2^F/+^; Myt3^F/F^; Pdx1^Cre^*. These mice were euglycemic and fertile up to 8-month after birth, allowing the above crossing. *6F* littermates were used as controls.

For *Hsp* OE, *TetO^3H^* mice (males and females) were crossed with *Rip^rTTA^* mice (females and males). The day of vaginal plug appearance was counted as embryonic day 0.5 (E0.5). To induce *Hsp* OE, pregnant mice were put on water supply with 200 microgram/ml Dox from E16.5 *ad lib* until the day of tissue collection. For initial Hsp OE characterization, via eGFP expression, all five lines were used. Two were euthanized because of their low (<10%) penetrance of eGFP production in β cells. Three had similar portions of β cells expressing eGFP, so that one was randomly chosen for all studies presented here. For testing the potential silencing of *Rip-rTTA* or *TetO^3H^* transgenes in adult ages, isolated islets from *Hsp-OE; Rip-rTTA* mice were treated with Dox at 2-month of age without prior exposure. For *ATF4* OE, a similar crossing scheme as above was used, except that the *Rosa26-ATF4^LoxTG^* and *Ins1^Cre^* mouse lines were used and no Dox was introduced. *Rosa26-ATF4^LoxTG/+^* and *Ins1^Cre^* littermates were all included as controls.

### Blood glucose, plasma hormone assays, and insulin tolerance test (ITT)

Fasting glucose levels were read via tail snip after 6-hour (before one month old) or overnight (after one month old) fasting. For plasma insulin assay, retro-orbital blood collection was used. For ITT, mice were fasted for 4 hours. Insulin was injected at 1 unit/kg. Blood glucose was then measured. The blood glucose levels were read with a NovaMax^plus^ meter using blood from tail tip.

### Pancreatic islet isolation

Pancreata were directly digested (for P1, P7, and P14 pancreata) (Huang and Gu, 2017) or perfused (for pancreata older than 2 weeks) with 0.5 mg/ml Type IV collagenase dissolved in Hanks Balanced Salt Solution (HBSS) with Ca^2+^/Mg^2+^. After digestion at 37 °C, lysates were washed in RPMI 1066 with 5.6 mM glucose and 10% fetal bovine serum (i.e. RPMI-FBS) 4 times. Islets were hand-picked in RPMI-FBS for downstream usage.

### Islet insulin secretion assays

For static insulin secretion, the % of total insulin secreted within a 45-minute window was measured unless noted. Hand-picked islets were allowed to recover in RPMI-FBS for 2 hours or overnight. Islets were washed twice with pre-warmed KRB solution (2.8 mM glucose, 102 mM NaCl, 5 mM KCl, 1.2 mM MgCl_2_, 2.7 mM CaCl_2_, 20 mM HEPES, 5 mM NaHCO_3_, and 10 mg/ml BSA, pH 7.4) and then incubated in KRB (37 °C) for one hour, washed with pre-warmed KRB once more. 10-15 islets were then transferred into each of the wells of 12-well plates with 1 ml pre-warmed KRB to start the secretion assays. For all assays, four or more mice of each genotype were used for islet isolation, with islets from two or more mice mixed and examined as 2-3 technical replicas. Insulin was measured with an Elisa kit from ALPCO following manufacturer protocol. Assays from human islets were performed with the same way, except the basal glucose was 3.3 mM.

For perifusion assays, hand-picked islets were incubated overnight in RPMI with 10 mM glucose for recovery. Islets were then placed in a 1-ml perifusion chamber, equilibrated in KRB with 5.6 mM glucose for 30 min and then challenged with 16.7 mM glucose, 5.6 mM glucose + 50 μM IBMX, and 5.6 mM glucose + 20 mM KCl. The perifusion fractions were collected in 3-min intervals at 1 ml/min flow rate and assayed for insulin by radioimmunoassay.

### Real-time PCR-based gene expression assays

RNA was prepared from hand-picked islets with TRIzol (Life Technologies) and a DNA free RNA^TM^ kit (Zymo Research). 20-100 ng total RNAs were then used for cDNA preparation and real-time PCR, utilizing SYBR green master mix of the Bio-Rad system, with oligos listed in Table S3. Note that the human *ATF4* and mouse *Atf4* coding sequences are conserved, making it possible to assay both cDNAs with a single pair of oligos (Table 3).

### Protein pulse labeling assays

Hand-picked islets were allowed to recover for ∼2 hours in RPMI-FBS media. Islets were then transferred to media with 1:4 mix of RPMI-1640: Cys /Met-depleted DMEM with 20 mM glucose and 10% FBS, supplemented with ^35^S-labeled Cys/Met. After four hours, labeled islets were incubated in RPMI-FBS for 15 minutes, washed in PBS, and lysed for gel-electrophoresis and quantification (aided with Image J). The number of cells from each islet sample was determined via real-time PCR to compare the relative copy number of genomic DNA. Lysates of same numbers of cells were loaded into each lane of the protein gels for comparison, followed by gel drying and film exposure.

### Immunolabeling, β-cell mass assays, and transmission electron microscopy

Antibody staining followed standard procedures. Briefly, pancreatic or other issues were dissected. They would be: 1) directly frozen, sectioned, and then fixed in 4% paraformaldehyde for 15 minutes followed by permeabilization and antibody staining with Ki67 or MafA antibodies; 2) fixed in 4% paraformaldehyde overnight at 4 °C, washed with PBS 3 times, frozen, and sectioned for transcription factor and Glut2 staining; 3) fixed in 4% paraformaldehyde overnight at 4 °C, washed with PBS 3 times, and prepared as paraffin sections. In this case, the tissue will be sequentially dehydrated with 45, 70, 80, 95, and 100% ethanol, and twice with xylene, and embedded in paraffin. For hormone staining, the paraffin block will be sectioned, rehydrated with the above xylene and ethanol solutions in reverse orders, and staining with antibodies. Small tissues (<0.5 mm in each dimension) can be directly fixed. Large pancreata (older than P14) were cut into 5-10 small pieces and fixed. Six and 20 μm thick sections were prepared for paraffin and frozen tissues, respectively. Sections were stained with antibodies diluted in basal solution (1X Phosphate Buffered Saline + 0.1% triton100 + 0.1% Tween-20 + 0.1% BSA + 0.05% donkey serum). Antibodies used were listed in Key Resource Table, with dilution from 1:500 to 1:5000, depending on the antibody used and amount of tissue on each slide (0.1 ml antibody mix for each slide). For imaging, laser-scanning microscopy (LSM) was used. When expression levels of a protein between samples were compared, slides/cells were processed side-by-side, and images were captured utilizing identical optical/electronic settings. For protein level quantification, LSM images taken under identical parameters were selected for Image J-based particle quantification. Double blind tests were used.

Transmission electron microscopy (TEM) followed established procedure. Briefly, islets were fixed in 2.5% glutaraldehyde in 0.1M cacodylate buffer (pH 7.4) overnight at room temperature. Islets were then washed and treated in 1% osmium tetroxide in 0.1M cacodylate buffer for 1 hour. Islets were then washed, embedded, thin-sectioned for imaging. For unbiased ER examination, image capture and quantification were performed under double-blind settings.

For quantifications of β-cell mass and replication index throughout the pancreas, the pancreatic block was sectioned at 20 μm intervals. One third to one fifth of all sections were labeled and scanned using Aperio ScanScope or an Olympus X51. Β-cell mass was then calculated based on the total pancreas weight and the percentage of tissues area that labeled for insulin.

### Βeta-cell mitosis and apoptosis assays

For mice younger than P10, frozen pancreatic sections were prepared for Ki67 staining and quantification. For older mice, multiple-day BrdU feeding via drinking water (50 ng/ml) were performed. The pancreata were recovered, sectioned, and stained for % of Ins+ cells that incorporated BrdU. Up to 5% of all islet sections were quantified in each mice. Alternatively, hand-picked islets were dissociated into single cells, spun onto glass slides, and stained for marker expression and counting. For apoptotic cell counting, similar scheme was used.

### Human islet culture

Hand-picked human islets (>95% purity) were cultured in CRML-1066 media with 10% FBS and 3.3 mM glucose overnight. For high glucose treatment, media were switched to 10% FBS with 3.3 or 20 mM glucose. For palmitate treatment media were switched to that containing 0.5% BSA or 0.6 mM palmitate with 5% FBS. Islets were cultured for ∼40 hours and assayed for GSIS and mRNA expression. For immunoassays, islets were washed 2X with PBS and dissociated into small cell clusters (single to ∼20-cell) and cytospun onto slides for after-fixation and staining/imaging.

### Gene expression in human EndoC-βH1 cells

Human EndoC-βH1 cells were grown in DMEM containing 5.6 mM glucose, 2% BSA, 50 μM 2-mercaptoethanol, 10 mM nicotinamide, 5.5 μg/mL transferrin, 6.7 ng/mL selenite, 100 units/mL penicillin, and 100 units/mL streptomycin (Ravassard et al., 2011). For siRNA transfection [with a mix of three individual siRNAs (20 nM each)], dissociated cells were incubated with siRNA (with RNAiMax) for 5 minutes before plating. Gene expression assays were performed three days after infection or transfection by directly lysing cells on plates for RNA assays or recovered, cyto-spun onto glass slides for protein expression assays. Human islets were purchased from IIDP or obtained from the Clinical Islet Transplantation Laboratory, University of Louisville, KY. Hand-picked islets were dissociated into single cells and Cyto-spun onto glass slides for antibody staining.

### Protein abundance assay at per cell levels

Stained cells (with immunofluorescence, dilution were marked on the key reagent Table) using tissue sections or dissociated islet cells that were cyto-spun onto glass slides. Mutant/OE and control slides were processed side by side. Confocal images were taken under identical parameters using non-saturating conditions. Image J was then used to quantify the fluorescence intensity. For Glut2 and insulin assays, the averaged fluorescence intensity within selected areas was used. For transcription factor assays, nuclei were circled and assayed for intensity. For background subtraction, areas outside the islets and inter-nuclear areas were assayed. Three to four pancreata were assayed, with three to ten representative microscopic areas assayed.

### Luciferase assays

The luciferase reporter construction followed routine molecular cloning process. Briefly, BAC clones carrying the *Hsp* genomic regions were purchased from Oakland Children’s Hospital. PCR were then used to amplify the genomic fragments indicated in Figure 5A and inserted into a vector containing firefly luciferase-eGFP fusion and a PolyA signal. The constructs were then sequenced to ensure correct sequences. Note that for unknown reasons we could not clone DNA fragments that contain a putative Myt1 and Myt2 binding motif ∼ 6 kilobase upstream of the *Dnajb1* transcription initiation site (Mall et al., 2017; Vasconcelos et al., 2016). The functional significance of this motif was therefore not tested.

For making firefly luciferase reporters with the 33-bp Myt1-binding sites of *Hspa1a,* a minimal CMV promoter was synthesized and ligated with the *Hspa1a* enhancer and the firefly luciferase reporter. The artificial Myt1ZF-VP16 constructs were made by PCR fragment ligation: the 4-zinc finger region was amplified with PCR, using a full length Myt1 cDNA as template; the VP-16 region was PCR-amplified with the rTTA cDNA as template. The oligos used were in Table S3. Luciferase assays followed established protocols. Briefly, HEK293T cells were transfected with Myt1 overexpressing or control plasmids, together with firefly luciferase reporters and a Renilla luciferase internal control. Cells were then lysed two days after transfection to assay the firefly and Renilla luciferase activity using a Dual Luciferase Assay kit (Perkin Elmer) following the manufacturer’s protocol.

### Putative Myt-binding site identification and ChIP-PCR assays

The putative Myt1 and Myt2 binding sites were identified by examining the supplementary tables in (Mall et al., 2017; Vasconcelos et al., 2016) based on ChIP-seq data and in silico search of the AAGTT motives recognized by the Myt TFs. ChIP assays used a Magna-CHIP^TM^ HiSens kit from Millipore, following manufacturers recommendation. Briefly, adult islets and acinar clusters were handpicked, fixed in 1% formaldehyde for 14 minutes and washed 2X with cold PBS. Islets were then frozen for storage. When ∼3000 islets were obtained, they were thawed and dounced into single cells in lysis buffer provided in the kit, and sonicated to ∼100-400 base pair fragments. Chromatin preparation from acinar cells followed the same process. The immunoprecipitation was done using 1 μg purified antibodies per 0.4 μg chromatin overnight. Oligos used in the following real-time PCR were listed in Table S4. Oligos from albumin promoter regions were used as controls.

### Sample size and statistical analysis

All experiments contained at least two biological replicas and two technical replicas, so that all assays had at least 4 independent experiments. Statistical analyses utilized standard Student’s *t*-test for pairwise comparisons or one-way ANOVA for comparing multiple groups of data points. A *p*-value of 0.05 or lower was considered significant.

### RNAseq Data analysis

The original RNA-seq data were deposited in ArrayExpress under ID code E-MTAB-2266 (https://www.ebi.ac.uk/arrayexpress/experiments/E-MTAB-2266/) and E-MTAB-6615 (https://www.ebi.ac.uk/arrayexpress/experiments/E-MTAB-6615/). The data were generated with purified β cells using *Mip-eGFP* expression as a surrogate for insulin production (Huang et al., 2018). After downloading the data, raw reads were processed and analyzed with TopHat and Cufflinks to determine the relative abundance of gene expression at each stages, reported as log2 transformed FPKM (Fragments Per Kilobase of transcript per Million mapped reads, Table S2)(Trapnell et al., 2012). To avoid excluding false negative genes, we included all genes with adjusted p-value smaller than 0.05. We also included other genes with a difference above 2-fold and a p-value smaller than 0.02.

For a full description of RePACT (regressing principle components for the alignment of continuous trajectory)-based human β-cell analysis, please refer to the star methods of (Fang et al., 2019). Briefly, we used Dropseq to generate massively parallel single cell transcriptome data (∼39,000 β cells, from 1,800 human islet). In RePACT, we first performed PCA to reduce the dimension of transcriptome data. Next, we used regression analysis to draw two optimal trajectory lines reflecting the T2D-relevant variation. In this study, we used the top 10 PCs as predictors. The numeric projection of each cell on the T2D trajectory (T2D index) served as a measurement of the degree to which the cell has transformed during disease development. We then binned the cells into a number of pseudo-states according to the index values. By comparing cells from different pseudo-states, RePACT greatly improved the statistical power to identify gene signatures for T2D status in α or β cell.

## KEY RESOURCES TABLE

| **REAGENT or RESOURCE** | **SOURCE** | **IDENTIFIER** |
| --- | --- | --- |
| **Antibodies (with fold of dilution indicted)** |  |  |
| 488-donkey anti Guinea Pig (1:1000) | Jackson Immunoresearch | 706-545-148<br>RRID:AB_2340472 |
| 488-donkey anti goat igG (1:1000) | Jackson Immunoresearch | 705-545-147<br>RRID:AB_2336933 |
| Alexa Fluor® 647 AffiniPure Donkey Anti-Guinea Pig IgG (H+L) (1:1000) | Jackson Immunoresearch | 706-605-148<br>RRID:AB_2340476 |
| Alexa Fluor® 488 AffiniPure Donkey Anti-Rabbit IgG (H+L) (1:1000) | Jackson Immunoresearch | 711-545-152<br>RRID:AB_2313584 |
| Alexa Fluor® 647 AffiniPure Donkey Anti-Rabbit (1:1000) | Jackson Immunoresearch | 711-605-152<br>RRID:AB_2492288 |
| Alexa Fluor® 647 AffiniPure Donkey Anti-Mouse IgG (H+L) (1:1000) | Jackson Immunoresearch | 715-605-150<br>RRID:AB_2340862 |
| Alexa Fluor® 594 AffiniPure Donkey Anti-Mouse (1:1000) | Jackson Immunoresearch | 715-585-150<br>RRID:AB_2340854 |
| Alexa Fluor® 594 AffiniPure Donkey Anti-Goat IgG (H+L) (1:1000) | Jackson Immunoresearch | 705-585-003<br>RRID:AB_2340432 |
| Alexa 647 AffiniPure Rabbit Anti-Syrian Hamster IgG (H+L) (1:1000) | Jackson Immunoresearch | 307-605-003<br>RRID:AB_2339601 |
| Rat anti-BrdU | ABcam | Ab-6326<br>RRID:AB_305426 |
| biotin anti Rat (1:1000) | Jackson Immunoresearch | 712-065-151<br>(RRID: N/A) |
| Cy3-Donkey anti-mouse (1:1000) | Jackson Immunoreserach | 715-165-150<br>RRID:AB_2340813 |
| Cy3-streptavidin (1:1000) | Vector laboroatories | SA-1300 (RRID: N/A) |
| Cy5- streptavidin (1:1000) | Vector laboroatories | SA-1500 (RRID: N/A) |
| FITC anti Rabbit (1:1000) | Jackson Immunoresearch | 711-095-152<br>RRID:AB_2315776 |
| Goat anti-Pdx1 (1:2000) | ABcam | AB47383<br>RRID:AB_2162359 |
| Goat anti-insulin (1:500) | Santa Cruz | sc-7839<br>RRID:AB_2296108 |
| Guinea pig anti insulin (1:1000) | Dako | A0564<br>RRID: N/A |
| Rabbit anti-Myt1 (1:1000) | This paper | N/A |
| Mouse anti-glucagon (1:5000) | Millipore | MabN238<br>RRID:NA |
| Rabbit anti-Pdx1 (1:5000) | ABcam | AB47267<br>RRID:AB_777179 |
| Rabbit anti-Nkx6.1 (1:5000) | Gift of P. Serup | NA |
| Rabbit anti-MafB (1:1000) | Gift of R. Stein | NA |
| Rabbit anti-Cleaved Caspase 3 (1:500) | SYSY | #105173<br>RRID:AB_887838 |
| Rabbit anti-glucagon (1:100) | ABcam | AB92517<br>RRID:AB_10561971 |
| Rabbit anti-Ki67(1:500) | Abcam | Ab 15580<br>RRID:AB_443209 |
| Guinea pig anti-Myt2 | This lab | NA |
| Rat anti-Myt3 | This lab | NA |
| **Bacterial and Virus Strains** |  |  |
| Bl-21 | Vanderbilt MPB Core | NA |
| DH5-alpha | Vanderbilt MPB Core | NA |
| Xl-1 | Vanderbilt MPB Core | NA |
| **Biological Samples** |  |  |
| Bovine serum albumin | Sigma | A9418 |
| Donkey Serum | Jackson<br>Immunoresearch | 017-000-121<br>RRID:AB_2337258 |
| Fetal Bovine Serum | Atlanta Biologicals | S10250 |
| **Chemicals, Peptides, and other key reagents** |  |  |
| Collagenase from Clostridium histolyticum | Sigma | C5138 |
| DMSO | Sigma | D2650 |
| Doxycycline | Sigma | D9891 |
| dapi | Sigma | D9542 |
| Adenosine, periodate oxidized | Sigma | 7154 |
| paraformaldehyde | Sigma | P6148 |
| Millicell | Millipore | PIHA01250 |
| Trypsin | Sigma | C5138-5G |
| SuperScript III Reverse Transcriptase | Thermo | 18080085 |
| RNaseOUT Recombinant Ribonuclease Inhibitor | Thermo | 10777019 |
| RNaseOUT Recombinant Ribonuclease Inhibitor | Thermo | 10777019 |
| Tn5 enzyme for methylome DNA tagmentation | Home made |  |
| T4 DNA Polymerase | NEB | M0203S |
| Ampligase | Lucigen | A3210K |
| **Critical Commercial Assays** |  |  |
| High capacity cDNA synthesis | Applied Biosystems | 4368814 |
| SYBER Green qpcr mix | Biorad | 1705060 |

## Supplementary Tables

**Table S1:** Summary of quantification data used in figures.

**Table S2:** List of genes differentially expressed between P1 control and *6F; Pdx1Cre* Mip-eGFP^+^ β cells. Fold change (LogFC), p-value, adjusted p-value with Benjamini and Hochberg (BH) method, and gene expression levels in three controls and three mutants (reads per kilo-base per million reads, log2 transformed) were listed.

**Table S3:** Expression trajectory of several stress genes in human β cells during T2D development.

**Table S4:** List of oligo nucleotides used for genotyping and PCR.

## Supplementary Figure Legends

**Figure S1:**
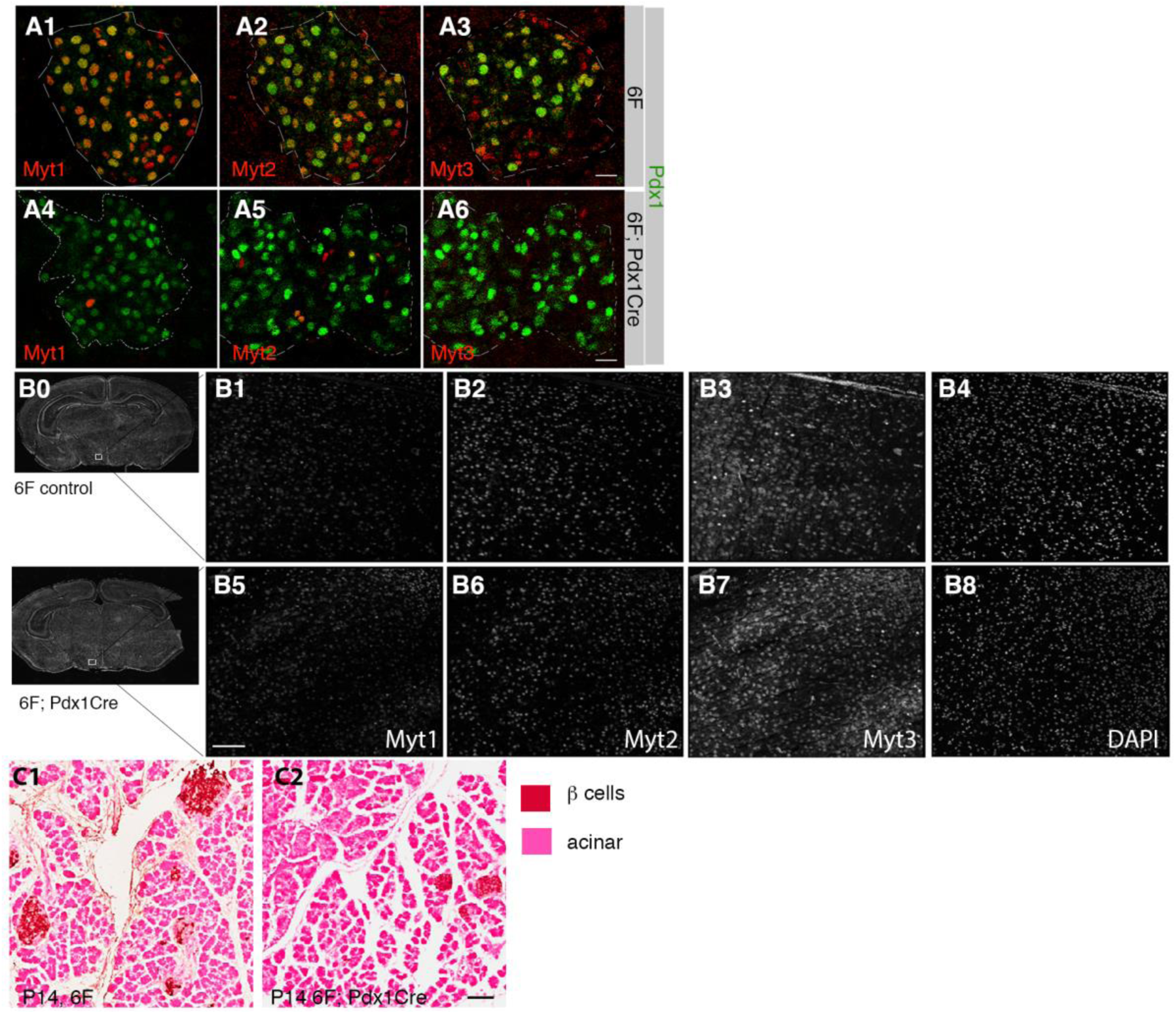
Quality control tests in *Myt* triple mutant mice. Related to Figure 1. Mice used here were derived from inter-cross of *Myt1^F/+^; Myt2^F/+^; Myt3^F/+^; Pdx1^Cre^* mice or between *6F* and *Myt1^F/+^; Myt2^F/F^; Myt3^F/F^; Pdx1^Cre^* or *Myt1^F/F^; Myt2^F/+^; Myt3^F/F^; Pdx1^Cre^* mice. The day of birth was counted as P1. For all quantification panels, the *p*-values [type 2, 2-tailed t-test except in G (ANOVA)] were marked on top of each assay. Error bars, SEM. “n” denotes the number of mice used. (A) Myt TF deletion efficacy in *6F; Pdx1^Cre^* islets, with a *6F* control presented (P14). Pdx1 signals (green) were used to identify the location of islets. Scale bars = 20 μm. (B) Myt TF detection in the hypothalamus regions of P14 *6F* and *6F; Pdx1^Cre^* mice, labeled for each Myt TF and DAPI. Scale bars = 50 μm. (C) The morphology of pancreatic exocrine tissues in P14 *6F* and *6F; Pdx1^Cre^* mice. Islets were labeled with HRP signals (brown). All cells were visualized by Eosin counter staining (pink). Scale bars = 100 μm.

**Figure S2:**
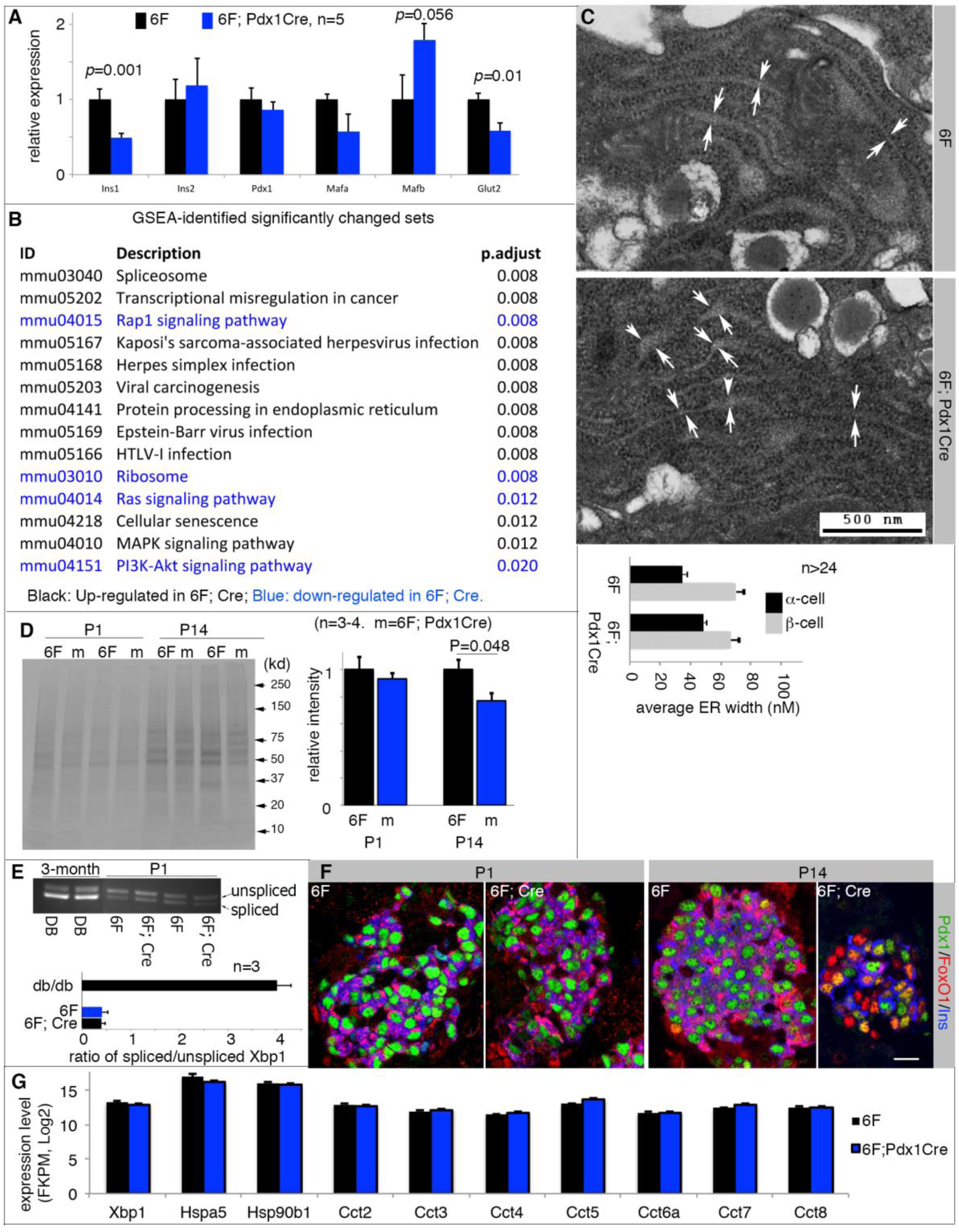
Myt TF mutant β cells lacked notable cellular stress before phenotype development. Related to Figures 2 and 3. Mice used here were derived from crosses between *6F* and *Myt1^F/+^; Myt2^F/F^; Myt3^F/F^; Pdx1^Cre^* or *Myt1^F/F^; Myt2^F/+^; Myt3^F/F^; Pdx1^Cre^* mice. The day of birth was counted as P1. For all quantification panels, the *p*-values (type 2, 2-tailed t-test) smaller than 0.05 were marked on top of corresponding assays. Error bars, SEM. “n”, the number of independent islet preparations used, except that in C, where “n” refers to the number of β cells counted. (A) Real-time RT-PCR assays of gene expression from P7 hand-picked islets. (B) Pathways differentially expressed in P1 *6F* and *6F; Pdx1^Cre^* β cells, identified via Gene Set Enrichment Assays. Pathways with adjusted *p* value ≤ 0.02 were listed. (C) ER-width in P1 β cells assayed via transmission electron microscopy. The presented quantification data were from double blind tests, counting the distance between two membranes of ER (distance between paired arrows). In the quantification, α cells were included as controls as well. (D) Radio-labeling (^35^S) and quantification of newly made proteins in *6F* and *6F; Pdx1^Cre^* islets within a four-hour window. The presented quantification data were from Image J-aided assays from exposed protein gels. (E) *Xbp1* mRNA spicing assays. Islets from 3-month old *db/db* mice were used as positive control, showing increased *Xbp1* splicing. The ratios between spliced (lower bands) and unspliced (upper bands) bands were presented (*6F*; *Cre*=*6F; Pdx1^Cre^*). (F) FoxO1 activation in *6F; Pdx1^Cre^* β cells. Pdx1 signals (green) were used to identify the β cells. Note the presence of yellow nuclei (FoxO1 activation) only in the *6F; Pdx1^Cre^* sample at P14. Scale bars = 20 μm. (G) Gene expression of several stress markers assayed via RNAseq, presented as Log2 (fragments per kb per million reads).

**Figure S3:**
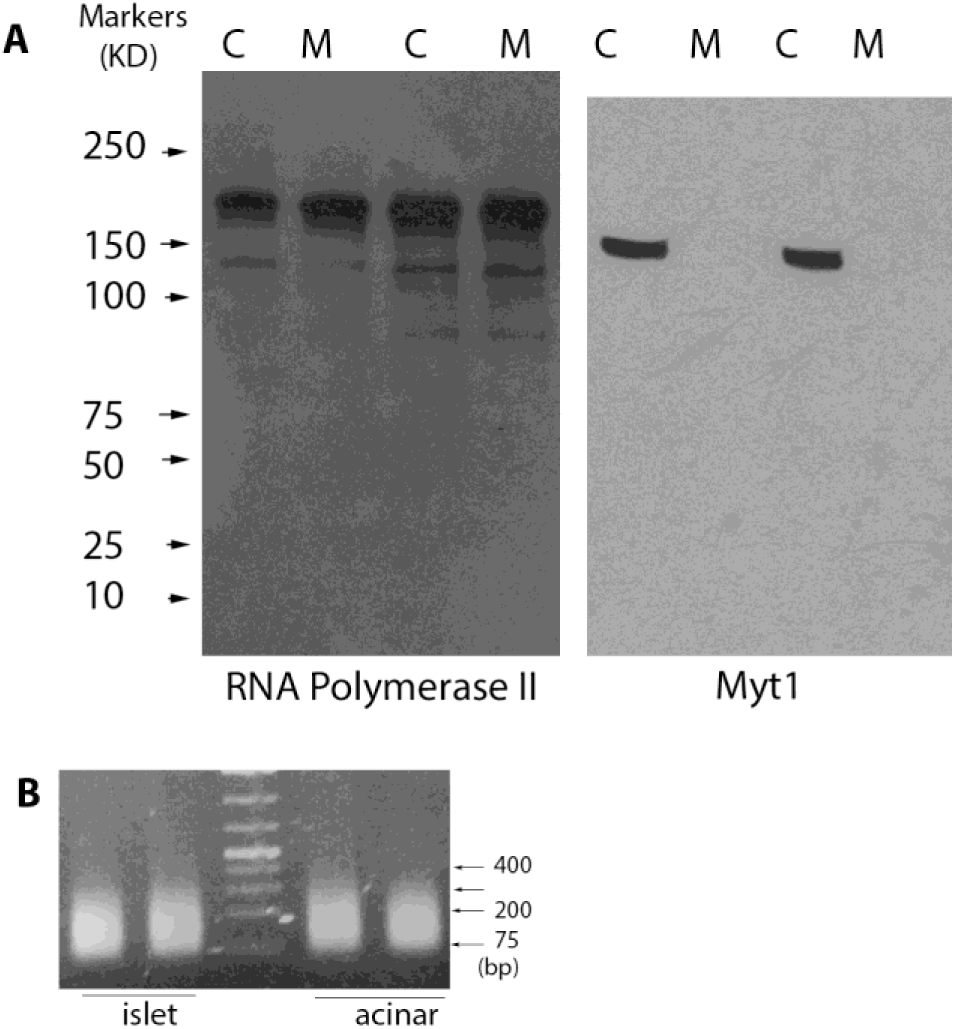
Myt1 CHIP assay controls. Related to Figure 3. (A) Western blot to test the specificity of the Myt1 antibodies used in CHIP-PCR (“C”, *6F* control islets. M, *6F; Pdx1^Cre^* mutant islets). An RNA polymerase II antibody was used as a positive loading/transfer control. (B) Examples of fragmented chromatin from adult islet and acinar cells.

**Figure S4.**
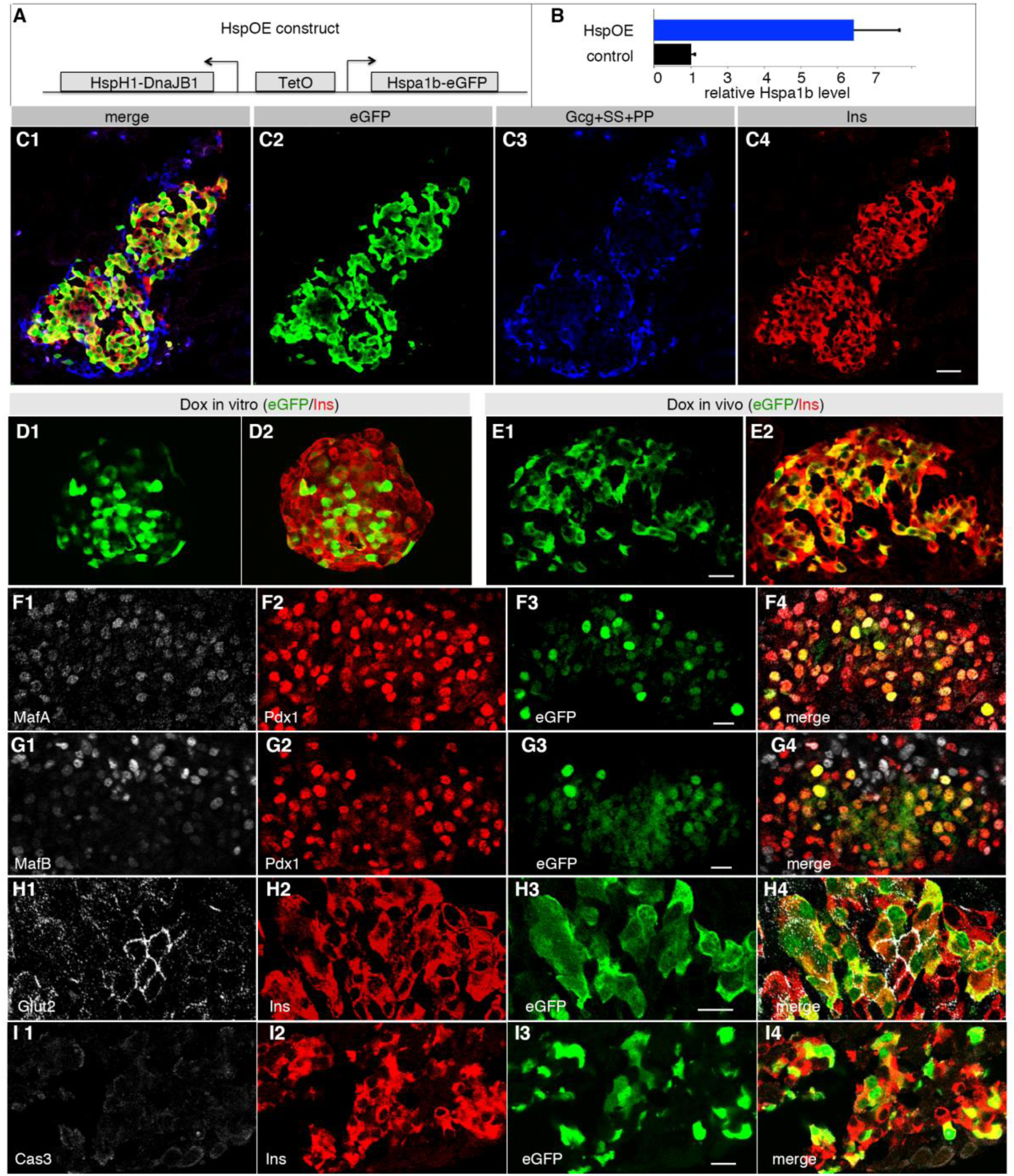
The derivation and some characterization of *Hsp-*overexpressing islet cells. Related to Figure 4. For inducing OE, *TetO^3H^* and *Rip^rTTA^* mice were crossed. Dox was administered in the drinking water of pregnant females starting from E16.5 (16^th^ day after the observation of vaginal plug). Dox application continued to the date of tissue collection. (A) The structure of *Hsp-OE* (*TetO^3H^*) transgene. Note that two mRNA will be transcribed under the control of a bi-directional TetO promoter. Each mRNA will produce two proteins, with T2A peptide included between HspH1 and DnaJB1 or Hspa1b and eGFP. (B) Level of *Hspa1b* OE in P2 *Hsp-OE* islets (*p*=0.01). Results were from four islet preps. Error bars, SEM. (C) Restricted transgene expression, by virtue of monitoring eGFP expression, in β cells of *Hsp-OE* islets (P4). Scale bars = 20 μm. (D, E) Hsp-OE in 8-week old islets (E) or mice (F) in animals without prior exposure to Dox but treated with Dox for two days. Islets were isolated from mice and treated with 5 ng/ml Dox for two days. For *in vivo* activation, two-month old mice were intra-peritoneally injected with 10 μg Dox. (F-I) Production of several β cell markers and Caspase 3 (Cas3) activation in P10 *Hsp-OE* islets. Scale bars = 20 μm. Single and merged channels were presented.

**Figure S5.**
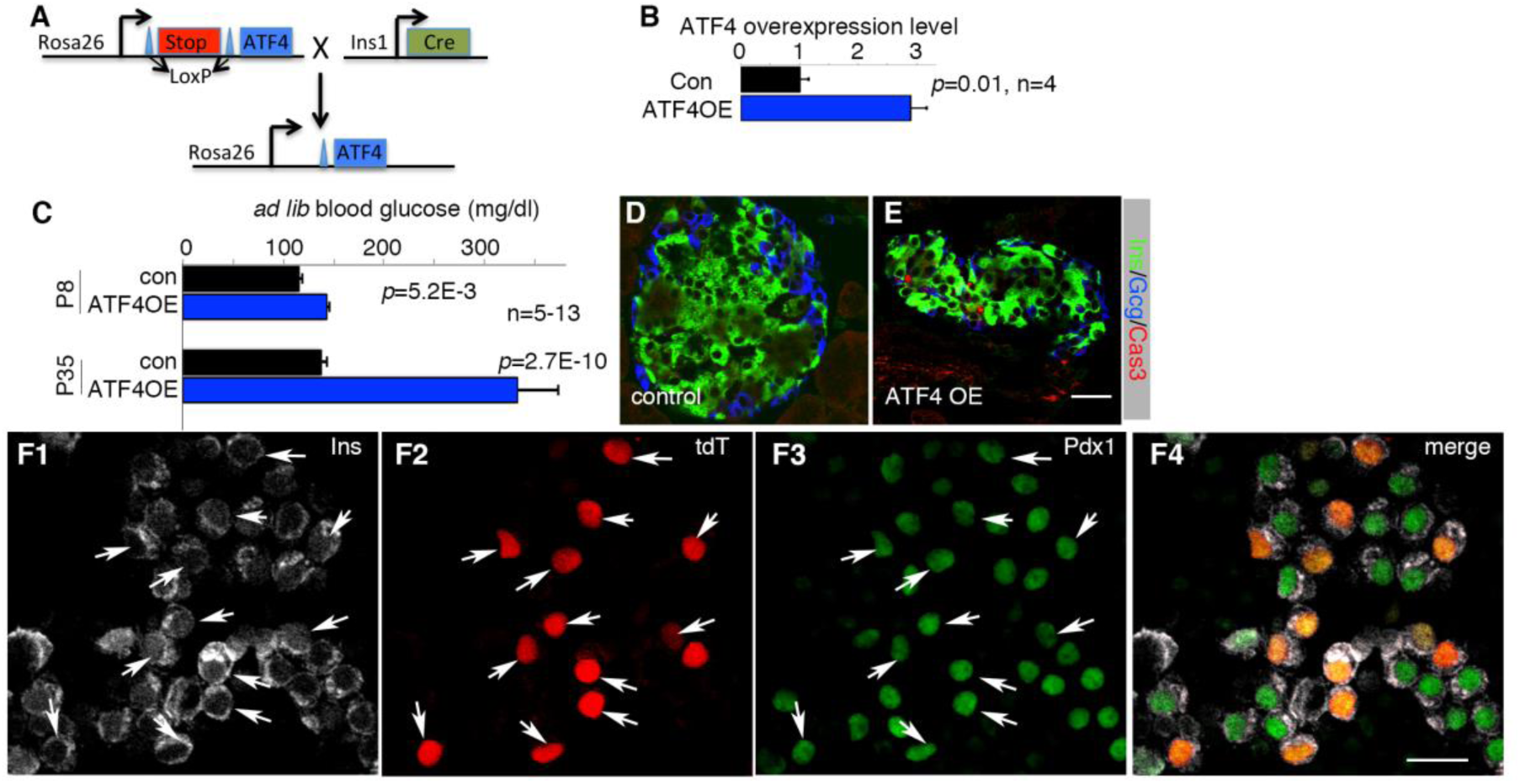
*ATF4 overexpression (OE) in mouse β cells*. Related to Figure 5. (A) The cross scheme used to activate *ATF4* OE in β cells. (B) Level of *ATF4* OE in P2 *ATF4-OE* islets. Results were from four islet preps. The P-values (type 2, 2-tailed t-test) were included in the panel. Error bars, SEM. (C) *Ad lib* blood glucose in control and *ATF4-OE* mice at two stages, before and after weaning. The controls include wild-type, *Ins1^Cre^*, and *Rosa26-ATF4^LoxTG^* mice, both males and females. The P-values (type 2, 2-tailed t-test) were included in the panel. Error bars, SEM. (D, E) An example of representative Cas3 activation in one-month old *ATF4-OE* islet. The few red dots in the ATF4OE panels are likely blood cells, which localize in the interstitial space between hormone-expressing cells. Scale bar = 20 μm. (F) Insulin and Pdx1 production in P35 control and tdTomato (tdT)-expressing β cells. tdT activation was achieved ∼E16.5, using a *Pdx1^CreER^*-based activation in an *Ai9* Cre-reporter mouse line. Arrows point to the tdT+ β cells. Compare the insulin (F1) and Pdx1(F3) signal intensities of tdT+ and tdT-cells. Scale bar = 20 μm.

**Figure S6.**
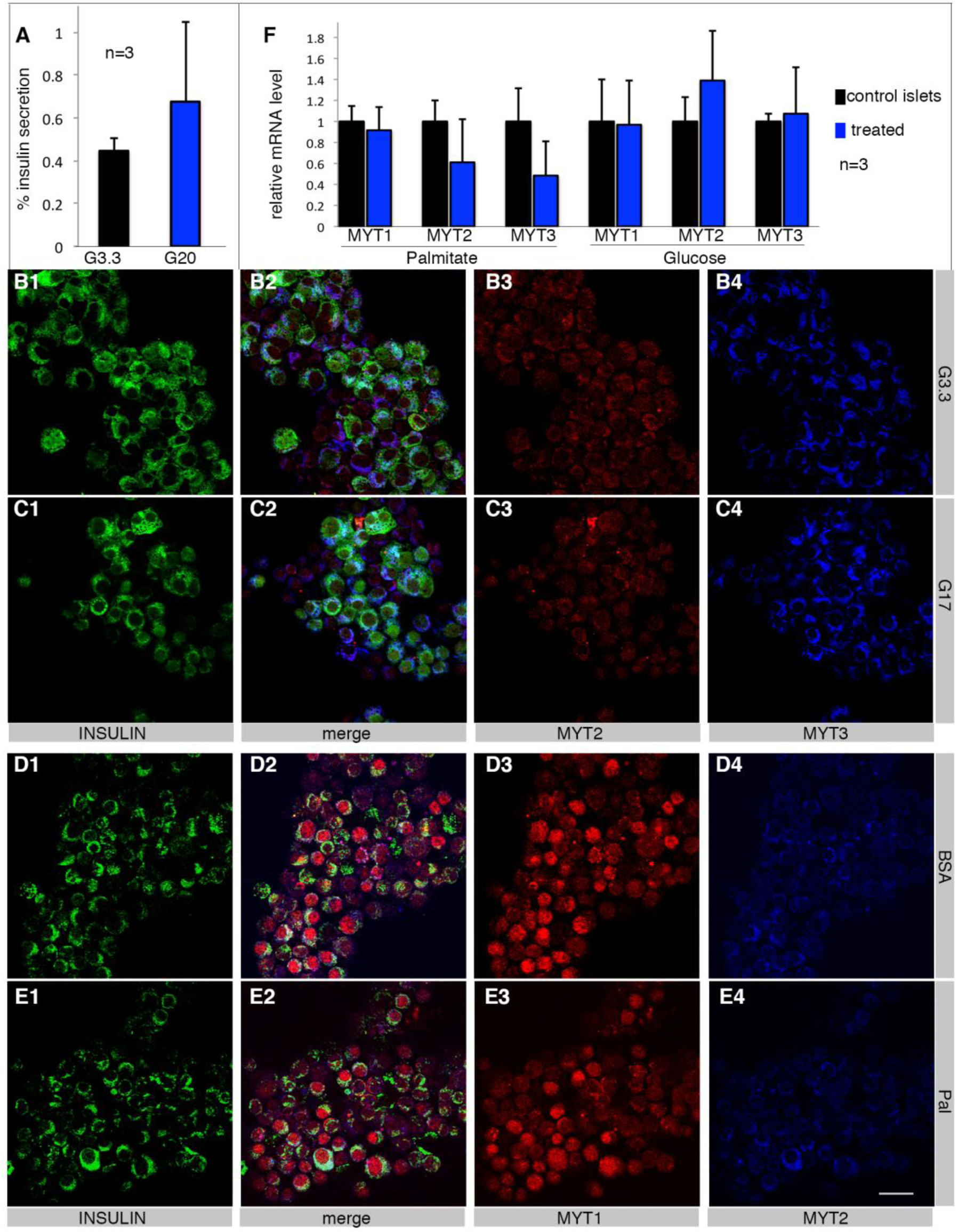
*MYT expression in human islet/β cells*. Related to Figure 6. (A) GSIS results of one batch of representative T2D islets (*p*=0.26). Shown are insulin secretion within a 30-minute stimulation window. (B, C) MYT2 and MYT3 in islet β cells (identified by insulin production) treated with 17 mM glucose for ∼40 hours. Starting materials were functional human islets. (D, E) MYT1 and MYT2 in islet β cells (identified by insulin production) treated with 0.6 mM palmitate for ∼40 hours. Starting materials were functional human islets. Scale bar = 20 μm, applicable to all images. (F) RT-PCR assays in human islets treated with G17 or palmitate for ∼40 hours. Whole islets were used for all assays. *P*>0.05 in all comparisons.

